# The vacuolar sugar transporter *Early Response to Dehydration 6-Like4* regulates fructose signaling and plant growth

**DOI:** 10.1101/2022.12.21.521376

**Authors:** Azkia Khan, Jintao Cheng, Anastasia Kitashova, Lisa Fürtauer, Thomas Nägele, Cristiana Picco, Joachim Scholz-Starke, Isabel Keller, H. Ekkehard Neuhaus, Benjamin Pommerrenig

## Abstract

Regulation of intracellular sugar homeostasis is maintained by regulation of activities of sugar import and export proteins residing at the tonoplast. We show here that the EARLY RESPONSE TO DEHYDRATION6-LIKE4 protein, being the closest homolog to the proton/glucose symporter ERDL6, resides in the vacuolar membrane. Gene expression and subcellular fractionation studies indicated that ERDL4 was involved in fructose allocation across the tonoplast. Overexpression of *ERDL4* increased total sugar levels in leaves, due to a concomitantly induced stimulation of *TST2* expression, coding for the major vacuolar sugar loader. This conclusion is supported by the finding that *tst1-2* knockout lines overexpressing *ERDL4* lack increased cellular sugar levels. ERDL4 activity contributing to the coordination of cellular sugar homeostasis is also indicated by two further observations. Firstly, *ERDL4* and *TST* genes exhibit an opposite regulation during a diurnal rhythm, and secondly, the *ERDL4* gene is markedly expressed during cold acclimation representing a situation in which TST activity needs to be upregulated. Moreover, *ERDL4*-overexpressing plants show larger size of rosettes and roots, a delayed flowering time and increased total seed yield. Consistently, *erdl4* knock-out plants show impaired cold acclimation and freezing tolerance along with reduced plant biomass. In summary, we show that modification of cytosolic fructose levels influences plant organ development and stress tolerance.

**One sentence summary:** The activity of the vacuolar sugar porter ERDL4 is important for balanced cytosolic monosaccharide homeostasis and influences plant growth and cold response in Arabidopsis

The author responsible for distribution of materials integral to the findings presented in this article in accordance with the policy described in the Instructions for Authors is: Benjamin Pommerrenig.

## Introduction

Sugars fulfill a wide number of functions in plants. They act as precursors for many anabolic reactions, allow starch and cellulose biosynthesis, serve as glycosyl donors for lipid and protein modification or provide the substrates for oxidative phosphorylation. In illuminated source leaves, sugar generation takes place in the cytosol of mesophyll cells, a process fueled by triose-phosphates exported from the chloroplast (Ruan, 2014). However, sucrose biosynthesis continues even in the dark phase, since stromal degradation of transitory starch provides maltose to the cytosol and the subsequent conversion of this disaccharide further drives sucrose biosynthesis (Smith and Zeeman, 2020).

Most vascular plants accumulate three major types of sugars in mature leaves, namely the disaccharide sucrose and the two monosaccharides glucose and fructose. These sugars are found in several cell compartments (Krueger *et al*., 2011; Fürtauer *et al*., 2019; Patzke *et al*., 2019) but most sugars are present in the vacuoles, representing the largest cell organelle (Martinoia *et al*., 2012). Interestingly, vacuolar sugar levels exhibit marked dynamic changes during the diurnal phase, but also in response to developmental demands or onset of various environmental stimuli (Smith and Stitt, 2007; Jänkänpää *et al*., 2012; Nägele and Heyer, 2013; Fürtauer *et al*., 2019). Accordingly, the vacuolar membrane harbors a range of transporters involved in either import or export of all major types of sugars (Hedrich *et al*., 2015). Thus, the abundances and finally activities of importers and exporters must be regulated. This regulation is realized by a complex network comprising various levels, like e.g., modified energization of the tonoplast, cell/organ specific expression patterns, altered gene expression in response to sugar availability or post-translationally via reversible phosphorylation (Cho *et al*., 2010; Wingenter *et al*., 2011; Schulze *et al*., 2012; Jung *et al*., 2015).

The individual sugar transporters so far demonstrated to locate to the vacuole belong either to the Monosaccharide Transporter (MST) group, or to the SUGARS WILL EVENTUALLY BE EXPORTED TRANSPORTER (SWEET) group (Eom *et al*., 2015; Pommerrenig *et al*., 2018). The Tonoplast Sugar Transporter (TST) and the VACUOLAR GLUCOSE TRANSPORTER1 (VGT1) have been the first tonoplast sugar transporters in Arabidopsis identified at the molecular level (Wormit *et al*., 2006; Aluri and Büttner, 2007). VGT1 seems to be specific for glucose transport, while TST was shown to accept monosaccharides and sucrose, and both carriers catalyze an electrogenic transport as they import their sugar substrates in counter-exchange to luminal protons (Aluri and Büttner, 2007; Schulz *et al*., 2011).

The export of luminal sugars from the vacuole is mediated by two types of carriers, namely those transporting sugars along their concentration gradient via facilitated diffusion, or those driven by the proton gradient in a sugar/proton symport mode. The EARLY-RESPONSIVE TO DEHYDRATION SIX (ERD6), and ERD6-LIKE (ERDL) 3 carriers (Kiyosue *et al*., 1998; Yamada *et al*., 2010) and the three SWEET proteins 2, 16, and 17 (Chardon *et al*., 2013; Klemens *et al*., 2013; Guo *et al*., 2014; Chen *et al*., 2015) catalyze glucose and/or fructose export. In contrast, the SUCROSE TRANSPORTER 4 (SUT4) and the ERDL6 protein are both proton-driven and allow for rapid export of sucrose or glucose, respectively (Poschet *et al*., 2011; Schulz *et al*., 2011).

From all known vacuolar sugar transporters, the TST type carriers represent the most in-depth analyzed proteins. In Arabidopsis, three TST isoforms exist: the *TST1* and *TST2* genes exhibit a tissue-specific expression pattern, while *TST3* is hardly expressed (Wormit *et al*., 2006). In contrast to *TST3, TST1* and *TST2* respond to stimuli like cold temperatures or sugar accumulation (Wormit *et al*., 2006). In storage taproots from sugar beet (*Beta vulgaris*), solely the isoform *BvTST2.1* is expressed and serves as a highly specific sucrose transporter responsible for the efficient sugar accumulation in this crop plant (Jung *et al*., 2015; Rodrigues *et al*., 2020). It is important to note that TST proteins do not only show a tissue-specific abundance but also exhibit altered phosphorylation patterns upon onset of cold temperatures (Schulze *et al*., 2012). This type of post-translational modification is catalyzed by a mitogen-activated protein 3-kinase family protein, named VIK1, and leads to increased transport activity (Wingenter *et al*., 2011). This process is fully in line with the observations that vacuolar sugar accumulation during cold temperatures is a key metabolic process induced to resist freezing (Pommerrenig *et al*., 2018) and that *tst1/2* double knock out mutants show strongly impaired cold-induced sugar accumulation when compared to wild types (Klemens *et al*., 2014).

Experiments on Arabidopsis or on false flax (*Camelina sativa*) lines constitutively overexpressing the *TST1* gene showed that corresponding mutants exhibit an increased source to sink transfer of sucrose promoting average seed size, lipid- and protein levels and the overall seed yield (Wingenter *et al*., 2010; Okooboh *et al*., 2022). The molecular basis for this effect is due to decreased cytosolic glucose levels in *TST1*-overexpressing plants, leading to a stimulation of photosynthesis and increased expression of the *SUC2* gene (Wingenter *et al*., 2010), encoding the major phloem sucrose loader (Truernit and Sauer, 1995; Lemoine *et al*., 2013). This stimulatory effect of increased TST activity on Arabidopsis seed size and total seed yield concurs with similar observations made on various crop plants. E.g., it has been recently shown that stimulated activity of TST leads to increased fruit sugar levels in strawberry- and melon fruits, as well as in apple and tomato fruits (Cheng *et al*., 2017; Zhu *et al*., 2021).

Interestingly, while stimulated fruit sugar levels in strawberry and melon fruits was achieved by a direct overexpression of the melon (*Curcumis melo*) *TST2* gene (Cheng *et al*., 2017), a stimulated TST activity in apple and tomato fruits was a consequence of overexpression of the apple (*Malus domestica*) gene *MdERDL6* (Zhu *et al*., 2021). The stimulated vacuolar glucose export in *MdERDL6-* overexpressing apple or tomato tissues lead to increased cytosolic glucose levels, which subsequently stimulate *TST2* expression. As a result of this process, the corresponding vacuoles are further loaded with sucrose finally leading to fruits with higher sugar concentrations when compared to wild types (Zhu *et al*., 2021). The observations made on leaves and fruits of apple and tomato mutants (Zhu *et al*., 2021) contrasts, however, findings made on Arabidopsis lines overexpressing the two closest *Md*ERDL6 homologs from either sugar beet (*Bv*IMP1) or Arabidopsis (*At*ERDL6). This is because overexpression of either *AtERDL6* or *BvIMP* leads to decreased, and not increased cellular sugar levels in leaves (Poschet *et al*., 2011; Klemens *et al*., 2014).

Thus, these contradictory observations make it mandatory to further analyze the coordination of sugar import and export across the tonoplast. To this end, we constitutively expressed the nearest ERDL6 homolog from Arabidopsis, *ERDL4* (Pommerrenig *et al*., 2018) in Arabidopsis. Interestingly, overexpression of *ERDL4* provokes increased source and sink tissue biomass, as revealed on rosettes, roots, and seed yield. In contrast, *erdl4* loss-of-function lines showed opposite characteristics. Further molecular and physiological data imply that fine control of a futile sugar transport cycle across the vacuolar membrane occurs in wild types and contributes to regulation of sugar allocation between source and sink tissues.

## Results

### Subcellular localization and functional characterization of the ERDL4 protein

In Arabidopsis ERDL4 belongs to the 53-member containing MST family of major facilitator superfamily proteins (Pommerrenig *et al*., 2018; Slavinski *et al*., 2021a). Within the MST family, ERDLs form a 19-member comprising subfamily making it the largest MST subgroup. Among ERDLs, ERDL4 has the highest homology to the vacuolar glucose/proton symporter ERDL6 sharing 92 % sequence identity (Poschet *et al*., 2010). Like all MSTs, ERDL4 possesses 12 putative transmembrane domains (TMD) (**Supplemental Figure S1**). The presence of sequences-PESPRWL- (between TMD six and seven), -DKAGRR- (between TMD eight and nine) and -PETKG- (after TMD twelve) constituting the highly conserved sugar transport motif (Sugar_tr; PF00083) suggested sugar transport properties (Henderson, 1991) (**Supplemental Figure S1 and Supplemental Information 1**).

Twelve ERDL members contain putative di-leucine-based motifs in their cytosolic N-termini (Yamada *et al*., 2010 and **Supplemental Information 1**) which constitute vacuolar targeting signals (Komarova *et al*., 2012; Wolfenstetter *et al*., 2012) and are required to target ERDL members ERD6 and ERDL3 to the tonoplast (Yamada *et al*., 2010). By contrast, this feature is absent from the ERDL4 protein sequence (**Supplemental Information 1**). To study the subcellular localization of ERDL4 experimentally, we expressed a construct containing the coding sequence (cds) of *ERDL4* and the *GREEN FLUORESCENT PROTEIN (GFP*) in frame under control of an *UBIQUITIN10-promotor* in tobacco mesophyll protoplasts. Recording of the GFP-derived fluorescence of the translated ERDL4-GFP with confocal microscopy revealed localization of the fusion protein at the tonoplast of intact (**Figure 1A–1D**) or lysed protoplasts (**Figure 1E, F**). These data showed that ERDL4 was targeted to the tonoplast independent of the presence of a dileucine motif.

**Figure 1.**
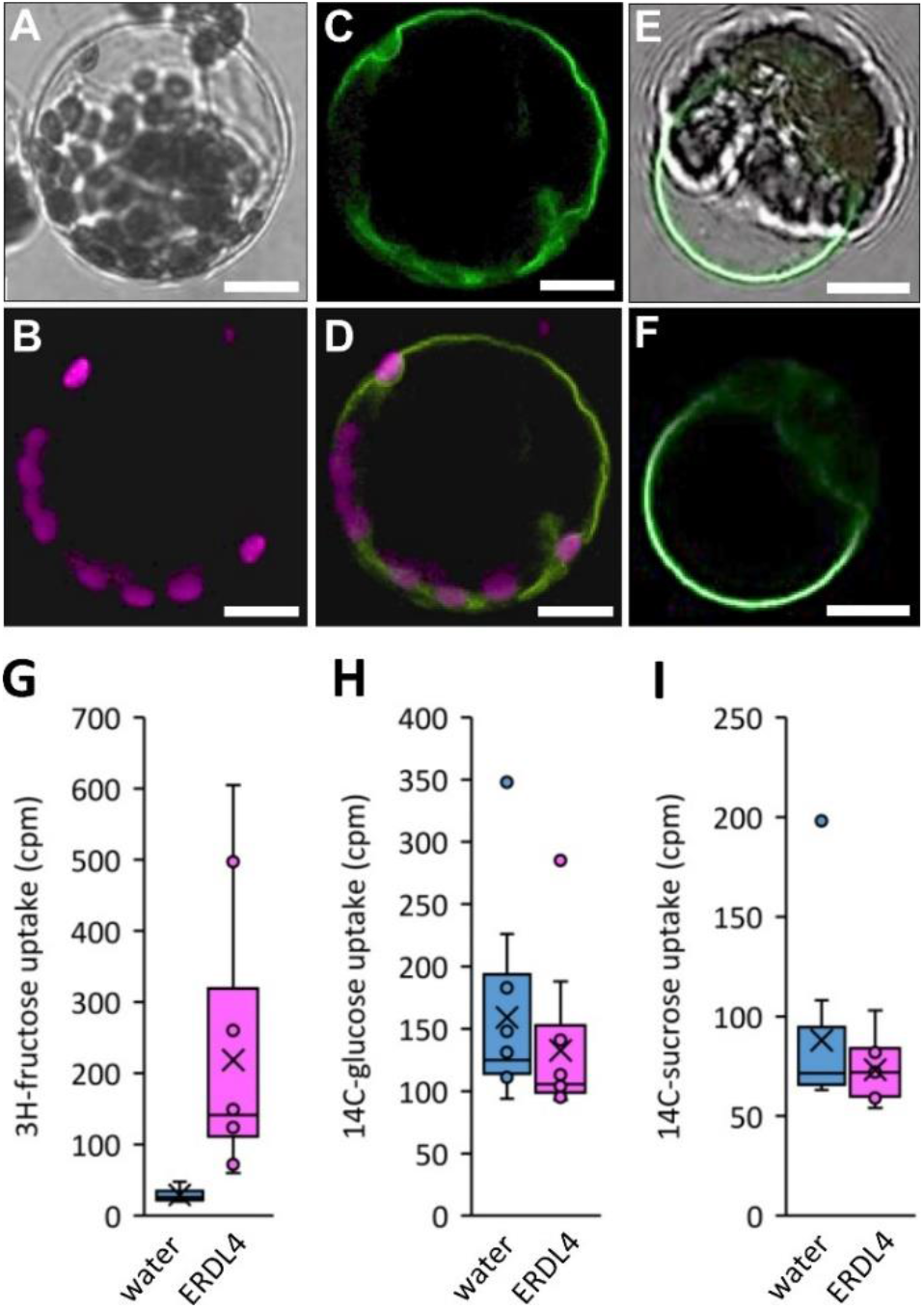
Subcellular localization of ERDL4-GFP fusion proteins in tobacco mesophyll protoplasts. **A-F)** Confocal images from tobacco mesophyll protoplasts transformed with an *UBQ10pro::ERDL4-GFP* construct. Scale bars are 10 μm **A-F)** Images of a tobacco protoplast expressing the ERDL4-GFP fusion. Different confocal channels of the same protoplast are shown **A)** Bright-field image **B)** Chlorophyll-derived auto-fluorescence indicating position of chloroplasts **C)** GFP-derived fluorescence indicating position of the ERDL4-GFP fusion protein **D)** overlay of chlorophyll and GFP fluorescence. **E and F)** images of a lysed protoplast expressing ERDL4-GFP **E)** GFP-derived fluorescence at the vacuolar membrane **F)** overlay picture of GFP fluorescence and bright field image **G-I)** Boxplots showing accumulation of radiolabeled sugars by oocytes injected with water (control) or *ERDL4*-cRNA. Center lines show the medians; box limits indicate the 25th and 75th percentiles; whiskers extend 1.5 times the interquartile range from the 25th and 75th percentiles, outliers are represented by dots; crosses represent sample means; n = 10 sample points. **G)** Uptake of ^3^H-fructose **H)** Uptake of ^14^C-glucose **I)** Uptake of ^14^C-sucrose. The p-values between water-injected and *ERDL4*-injected oocytes were 0.0046 (frc), 0.4082 (glc), and 0.3022 (suc) according to double-sided Students t-test.

It is noteworthy to mention that an ERDL4-GFP fusion transformed into the hexose transport-deficient yeast strain EBYVW-4000 (Wieczorke *et al*., 1999) did not target to yeast plasma membrane, as reported for the homologous proteins *Md*ERDL6 (Zhu *et al*., 2021) and *At*ERDL10 (Klemens *et al*., 2013), but to internal membrane structures (**Supplemental Figure S2**). To analyze substrate transport properties of ERDL4, we applied radiolabeled sugars - glucose, fructose, or sucrose - to *ERDL4-* expressing or water-injected (control) oocytes from *Xenopus laevis*. These analyses revealed that only ^3^H-fructose application increased the measured radioactive counts in the *ERDL4-*-injected oocytes significantly over the water-injected control oocytes (**Figure 1G–I**) and suggested that ERDL4 mediated the transport of fructose but not of glucose or sucrose.

### *ERDL4* is expressed in a diurnal manner

Dynamics of vacuolar sugar contents are linked to diurnal rhythm (Nägele et al., 2010) and vacuolar sugar transporters steer subcellular sugar compartmentation (Wormit *et al*., 2006; Poschet *et al*., 2010; Vu et al., 2020). We therefore checked whether *ERDL4* and other vacuolar sugar transporter genes were regulated in a diurnal manner. To this aim we grew wild-type plants in a 10 h light, 14 h dark regime (short days), harvested plant material in two-hour intervals, and analyzed expression of *ERDL4* as well as of *TST1* and *TST2* (Wormit *et al*., 2006; Schulz *et al*., 2011), encoding the two major transport proteins catalyzing sugar loading into the vacuolar lumen. In addition, we analyzed the expression of *ERDL6*, the closest homolog to ERDL4 catalyzing glucose/proton symport across the tonoplast (Poschet *et al*., 2010) (**Figure 2**).

**Figure 2.**
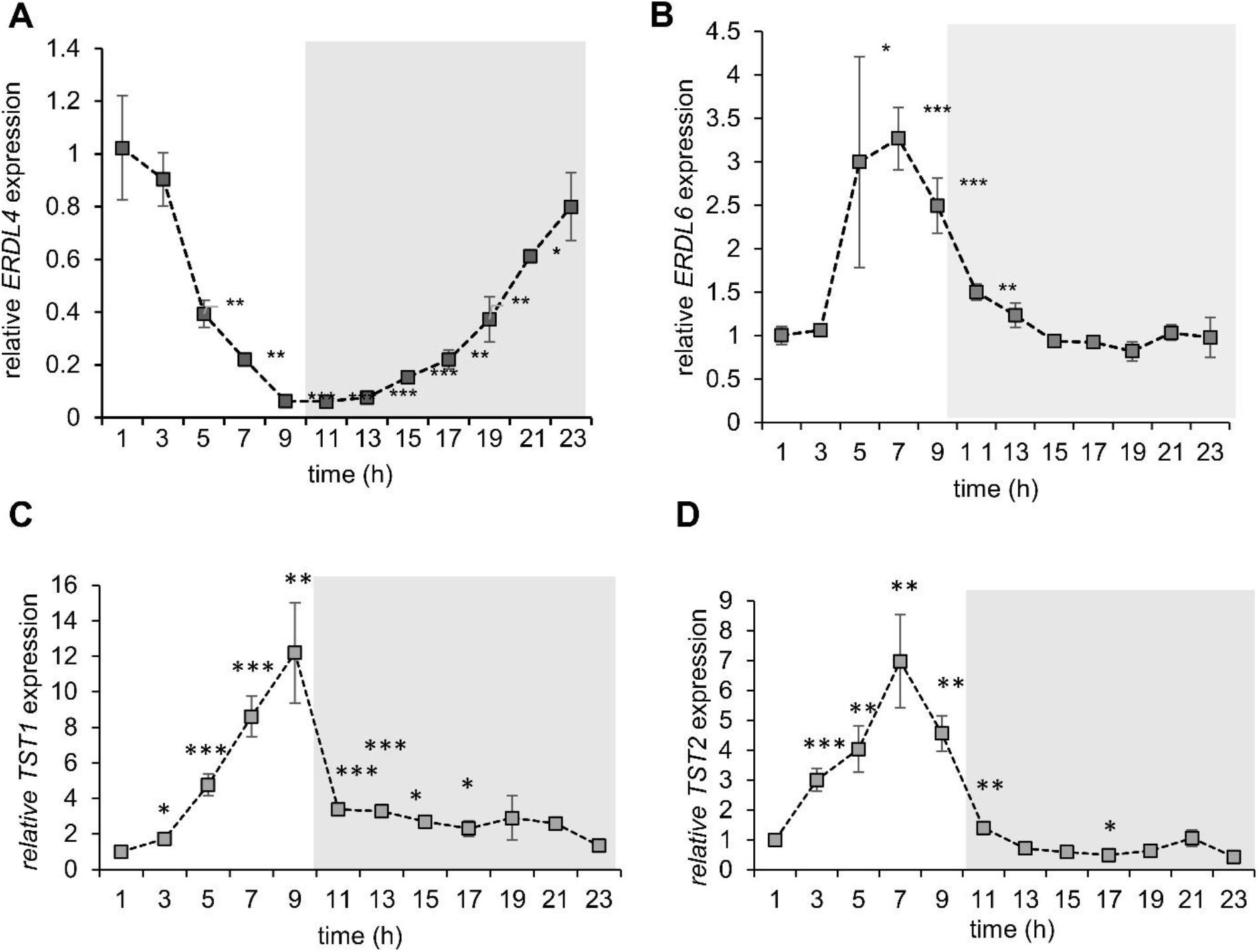
Diurnal expression of *ERDL4 (***A)**, *ERDL6* **(B)**, *TST1* **(C)**, and *TST2* **(D)**. Expression is shown as fold-change relative to the 1h-light value. Grey-shaded areas indicate nighttime (without illumination). Bars are mean ± SE values of n=3 replicates. Asterisks indicate significant changes to the expression at 1h according to double-sided Students *t*-test (* = p < 0.05; ** = p < 0.005; *** = p < 0.001).

With the beginning of the light phase *ERDL4* mRNA levels continuously decreased over the day (**Figure 2A**) reaching a minimum after nine hours in the light (= end of day) with about 16-fold less mRNA when compared to the first recorded value of the day. After the beginning of the dark phase *ERDL4* mRNA levels steadily increased until they regained their maximum from the start of the day after eleven hours in the dark (= end of night, EON). *ERDL6, TST1* and *TST2* genes on the contrary exhibited closely overlapping expression patterns and their mRNA levels sharply increased through the light phase peaking at seven to nine hours in the light phase. With the beginning of the dark period, the mRNA levels of these three genes rapidly decreased reaching the low expression levels from the start of the day again at the end of the night (**Figure 2B–D**). These results showed that expression patterns of the analyzed vacuolar sugar transporter genes were diurnally regulated and suggested that ERDL4 does not functionally overlap with TSTs or its closest homolog ERDL6.

### Isolation of *erdl4 T-DNA* insertion mutants and generation of *ERDL4*-overexpressing lines

To study effects of altered ERDL4-dependent vacuolar transport processes, we isolated *erdl4-* knockout mutants and generated *ERDL4-*overexpressing lines with decreased or increased ERDL4 activity, respectively. We isolated two independent *erdl4* mutant lines for our analyses, that carry T-DNA insertions either in the promotor region at −34 bp before the start-ATG (*SALK_11641; erdl4-1*) or at the beginning of the 14^th^ exon at +2281 bp (*GABI-KAT 600B02; erdl4-2*) (Alonso *et al*., 2003) (**Figure 3A**). RT-PCR based on leaf cDNA from the two mutants failed to amplify full-length *ERDL4-cDNA* from both *erdl4-1* and *erdl4-2* plants (**Figure 3B**). This characterized *erdl4-1* and *erdl4-2* as knockout mutants of ERDL4 and suggested that the insertion in *erdl4-2* mutants truncates the ERDL4 protein after the first three amino acids of the predicted ninth transmembrane domain (**Figure 3A, Supplemental Figure 1B**). In addition, we generated *ERDL4*-overexpressing lines expressing the *ERDL4-*cds under the transcriptional control of *the CaMV-35S-promoter* in Col-*0* background (*35S-ERDL4 #1* and *35S-ERDL4 #2*). RT-qPCR analyses demonstrated that the overexpression resulted in significantly higher (about tenfold for *35S-ERDL4 #1* and *#2*) amounts of *ERDL4* mRNA in these lines when compared to wild type (**Figure 3C)**.

**Figure 3.**
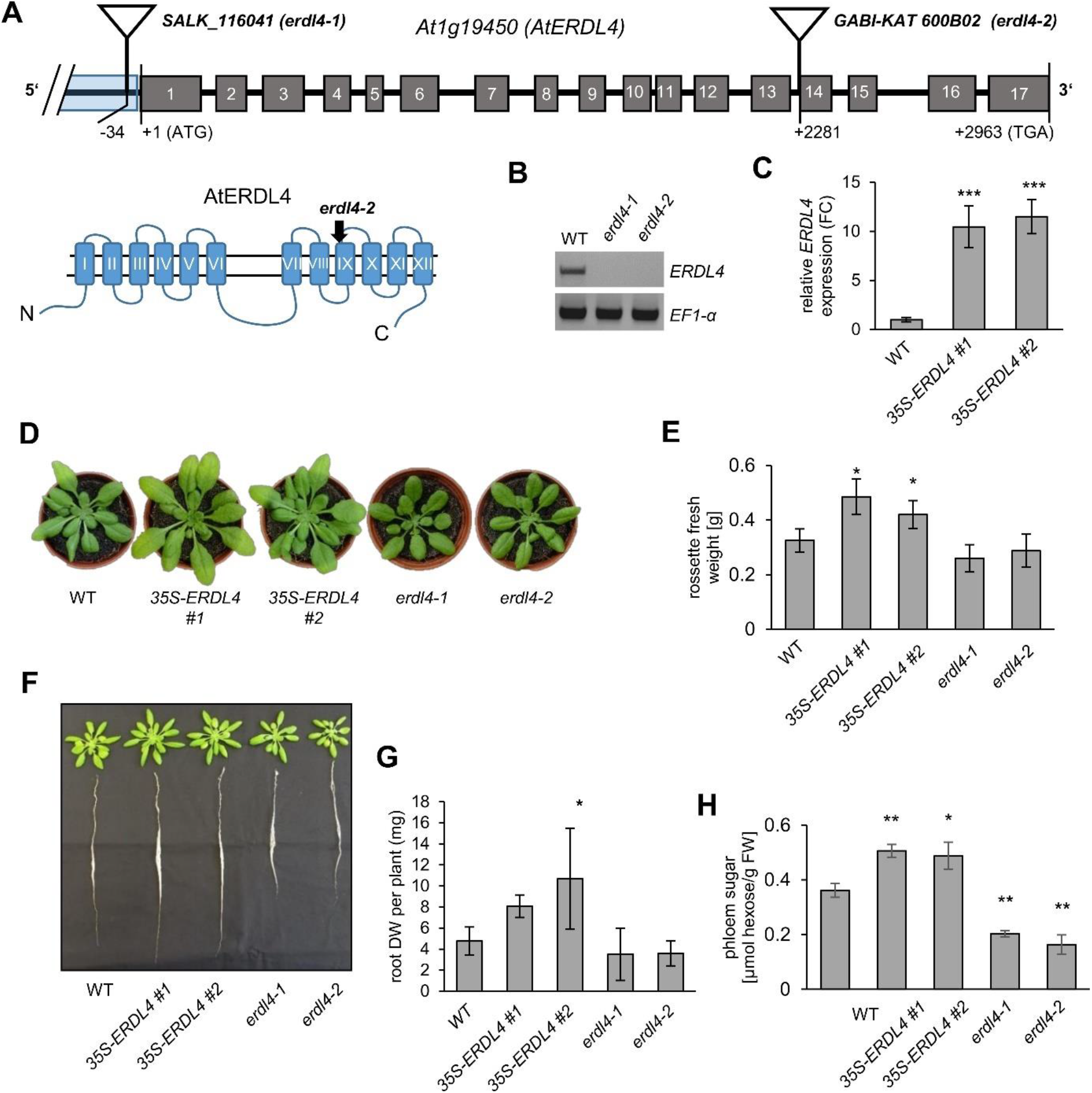
Characterization of *erdl4* T-DNA insertion mutants and *ERDL4*-overexpressing plants. **A)** Exon-intron structure of the *ERDL4* gene. Grey boxes with numbers indicate exons, black lines between boxes indicate intron sequences. Triangles mark T-DNA insertions in *erdl4-1* and *erdl4-2* mutant lines. Numbers below exons and introns indicate base pair positions of T-DNA insertions according to Alonso et al., 2003. The schematic below the gene shows the predicted secondary structure of the ERDL4 protein according to TMH-Pred. The black arrowhead marks the predicted truncation of the protein in *erdl4-2* mutants. **B)** RT-PCR analysis of *ERDL4* and *ELONGATIONFACTOR1-a (EF1-α*) expression in WT, *erdl4-1*, and *erdl4-2* mutants. **C)** Relative expression of *ERDL4* in WT and *35S-ERDL4* plants determined by RT-qPCR. Bars show mean values of n=4 homozygous plants ± SE. Asterisks indicate significant changes according to Students t-test in comparison to WT (*** p < 0.001). **D)** Phenotypes of four-week-old WT, *35S-ERDL4*, and *erdl4* knock-out plants grown in soil substrate under short day conditions. **E)** Quantification of shoot biomass on fresh weight basis of plants depicted in (D). Bars show mean values of n=12 independent plants ± SE. Asterisks indicate significant differences to WT according to Students *t*-test (* p < 0.05). **F)** Exemplary pictures of rosettes and roots from five-week-old WT, *35S-ERDL4*, and *erdl4* knock-out plants grown in hydroponic culture. **G)** Dry biomass of roots from plants depicted in (F). Bars show mean values of n=6 roots ± SE. **H)** sugar measured as hexose equivalents in phloem exudates collected from detached leaves of five-week-old WT, *35S-ERDL4*, and *erdl4* knockout plants. Bars are mean values of at least n=12 leaves ± SE. Asterisks indicate significant differences to WT according to Students *t*-test.

Interestingly, the altered *ERDL4* expression resulted in differential growth behavior of plants from the corresponding lines. Rosettes of five-week-old *35S-ERDL4 #1 and #2* plants grown on soil substrate were larger and had larger leaves in comparison to wild type plants of the same age (**Figure 3D**). These plants had about 50% (*35S-ERDL4 #1*) and 30% (*35S-ERDL4 #2*) more biomass on a freshweight basis when compared to wild type plants (**Figure 3E**). Rosettes of *erdl4.1* and *erdl.2* mutants from the same age showed decrease of their biomass to 80% (*erdl4-1*) or 90% (*erdl4-2*) of wild type values (**Figure 3E**). When grown in hydroponic culture, root biomass was increased in *35S-ERDL4 #1 and #2* plants and decreased in *erdl4-*knockout mutants when compared to WT plants (**Figure 3F, G**).

The altered *ERDL4* expression in the overexpressing or knock-out plants resulted in differential sugar contents of phloem sap collected from source leaves of plants from the different lines. While *35S-ERDL4* plants had between 30% and 40% higher sugar content of the phloem sap when compared to wild type, the levels were markedly reduced in *erdl4-1* and *erdl4-2* to about half of the wild-type content (**Figure 3H**). Together, these results characterized ERDL4 as an important factor for the growth of shoots and roots and indicated that ERDL4 activity influenced shoot to root transport of carbohydrates.

### Soluble sugar content of plants with altered ERDL4 activity

We analyzed whether altered activity of ERDL4 would influence whole leaf sugar content as well as intracellular compartmentation of sugars. Surprisingly, overexpression of *ERDL4* resulted in markedly increased accumulation of glucose in leaves of *35S-ERDL4* plants (**Figure 5A**). Glucose levels increased more than two-fold from 1.6 μmoles /g FW in WT to about 3.8 and 3.3 μmoles /g FW in *35S-ERDL4 #1* and *35S-ERDL4 #2* leaves, respectively (**Figure 5A**). In contrast, glucose was reduced to about 40% and 45% of wild type levels in leaves of *erdl4-1* and *erdl4-2* knock-out plants, respectively. Fructose contents were slightly, but not significantly increased in *35S-ERDL4* plants in comparison to the WT but reduced in *erdl4* knockouts (**Figure 5A**). Sucrose levels were not significantly altered between WT and other lines except for *35S-ERDL4 #1* plants, which exhibited about 70% higher sucrose levels when compared to WT (**Figure 5A**).

**Figure 5.**
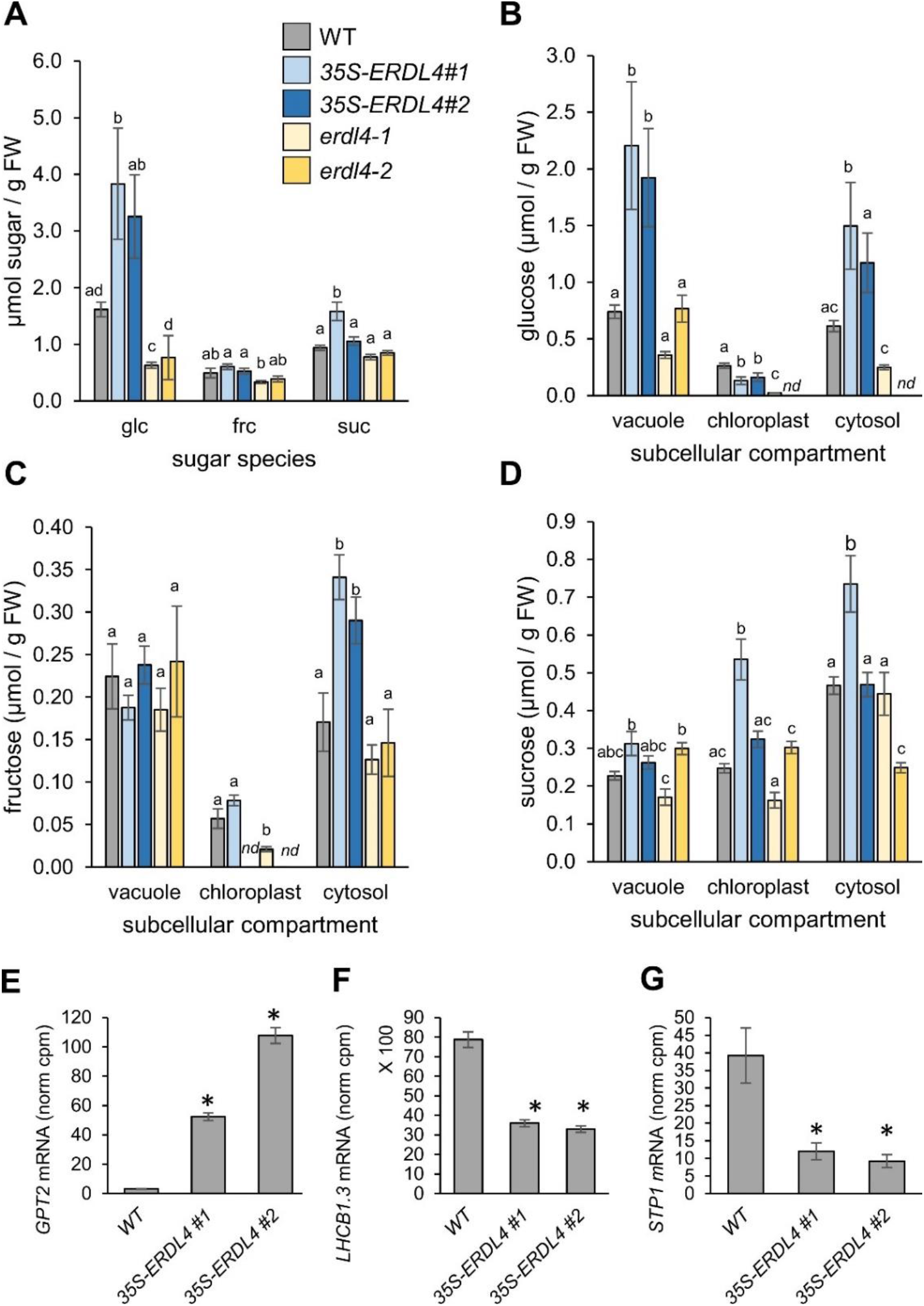
Total and subcellular sugar accumulation in WT, *ERDL4*-overexpressing and *erdl4* mutant plants. **(A)** Contents of glucose (glc), fructose (frc), sucrose (suc) in shoots harvested at midday (four hours in the light phase). **(B-D)** subcellular contents of glucose **(B)**, fructose **(C)**, sucrose **(D)** in vacuoles, chloroplast, and cytosolic fractions of shoots from WT, *ERDL4*-overexpressing and mutant plants as obtained by non-aqueous fractionation. Bars represent means from n=5 plants ± SE. Different letters above bars denote significant differences according to *One-Way* ANOVA with post-hoc Tukey test (p < 0.05). *nd* = no quantifiable amount detected. **E-G)** expression of sugar-responsive gens extracted from RNASeq-data in WT and *35S-ERDL4* plants. Data are means from n=3 biological replicates ± SE. Asterisks denote significant differences according to double-sided Students t-test (p < 0.05) **E)** expression of *GLC6-PHOSPHATE/PHOSPHATE TRANSLOCATOR2 (GPT2*) **F)** expression of *CHLOROPHYLL A/B BINDING PROTEIN1 (LHCB1.3; CAB1*) **G)** expression of *SUGAR TRANSPORT PROTEIN1 (STP1*).

To analyze the intracellular compartmentation of sugars, we conducted non-aqueous fractionation (NAF), allowing for quantification of sugars in the cellular compartments of vacuole, chloroplasts, and cytosol (Fürtauer et al., 2019; Patzke *et al*., 2019; Vu *et al*., 2020). We observed that altered expression of *ERDL4* lead to perturbations in the sugar homeostasis between the analyzed compartments (**Figure 5B to 5D**). Most prominent was the increase of glucose in the vacuolar and cytosolic but not in the chloroplast fractions of *35S-ERDL4* lines. Glucose increased about three-fold in vacuoles when compared to WT levels (**Figure 5B**). In the cytosol of *35S-ERDL4* lines this increase was about two-to 2.5-fold. Interestingly, the slight increase of total fructose (**Figure 5A**) in *35S-ERLD4* plants was not distributed proportionally over vacuole and cytosol (as observed for glucose) but was rather confined to the cytosolic fraction (**Figure 5C**). The distribution of sugars between compartments in *erdl4* knockouts did not significantly differ from those in WT except for *erdl4-1* plants which had significantly lower fructose levels in chloroplasts in comparison to the WT (**Figure 5C**). The elevated monosaccharide content in the cytosol of *35S-ERDL4* plants was further corroborated by altered expression of sugar-responsive marker genes in these lines (**Figure 5E–G**). Expression of *GLC6-PHOSPHA7E/PHOSPHA7E 7RANSLOCA7OR2* (*GP72*), *CHLOROPHYLL A/B BINDING PROTEIN1 (LHCB1.3; CAB1*), and *SUGAR 7RANSPOR7 PRO7EIN7 (S7P1*) has been shown to strongly mirror cytosolic glucose levels (Chen et al., 2019). While *GPT2* is sugar-induced, *CAB1* and *STP1* are sugar-repressed (Chen et al., 2019). The same pattern was observed for expression of these genes in the *35S-ERDL4* plants when compared to wild-type expression (**Figure 5E–G**). These data indicated that *ERDL4-*overexpressing plants had altered transport activities for both glucose and fructose at the vacuolar membrane. The observation that fructose levels of *35S-ERDL4* plants increased solely in the cytosol and not in the vacuole (as observed for glucose) suggested that ERDL4 mediated release of fructose from vacuoles to the cytosol.

### Differential gene expression in ERDL4-overexpressing plants overlaps with fructose rather than glucose induction

Overexpression of *ERDL4* resulted in elevated fructose and glucose levels in the cytosol and it is reasonable to check for consequences on global gene expression caused by the altered intracellular sugar distribution by means of RNASeq analysis. Having already seen that the elevated cytosolic sugar levels were transduced to the level of gene expression (**Figure 5E–G**), we were interested to check whether changes of the transcriptome of *ERDL4*-overexpressors were caused by increased cytosolic accumulation of glucose, or fructose or of both.

To this aim, control experiments were performed using excised Arabidopsis leaf discs that were incubated overnight in 0.5x MS buffer solution pH 5.7 supplemented with either 1% glucose, 1% fructose or 1% mannitol (as osmotic control). Under these conditions, the supplemented sugars or mannitol diffuse via the apoplasm to intact mesophyll cells where they are imported into the cytosol. Such flooding with pure sugar species has been shown to evoke sugar species-specific responses on the level of gene expression (Wormit *et al*., 2006; Luo *et al*., 2012). Genes which exhibit significant differential expression between different plant lines or between different treatments are named “differentially expressed genes” (DEGs). We identified DEGs between *ERDL4*-overexpressors and wild-type (WT), between fructose and mannitol incubation or between glucose and mannitol incubation by testing for significant differences of the expression of individual genes. Most genes regulated by one of the sugars were regulated by the other in the same manner as revealed by high correlation (R^2^ = 0.74) of the individual amplitudes of gene expression changes (as calculated as log2FC) between glucose- and fructose-regulated genes (**Supplemental Figure S3**). We checked if this was also the case in *35S-ERDL4* lines where both glucose and fructose levels were elevated in the cytosol at the same time.

We identified 130 DEGs (p < 0.001) that were common between *35S-ERDL4/WT* and the fructose/mannitol treatments and 155 DEGs between *35S-ERDL4/WT* and the glucose/mannitol treatments (**Figure 6A, B**). Correlation analysis of the 130 common DEGs from *ERDL4*-overexpressors and fructose and of the 155 common DEGs from *ERDL4*-overexpressors and glucose treatments revealed Pearson coefficients of *r* = 0.48 (*r^2^* = 0.23) and *r* = 0.15 (*r^2^* = 0.0225) for *ERDL4*-overexpressors and fructose treatment and *ERDL4-*-overexpressors and glucose treatment, respectively **(Figure 6C, D)**. This ten-times higher correlation between *ERDL4*-overexpressing- and fructose-dependent gene expression changes in comparison to *ERDL4*-overexpressing- and glucose-dependent gene expression changes indicated that fructose had a stronger effect on the transcriptional changes recorded in *ERDL4*-overexpressors in comparison to glucose and suggested a specific role of fructose in cytosolic sugar signaling.

**Figure 6.**
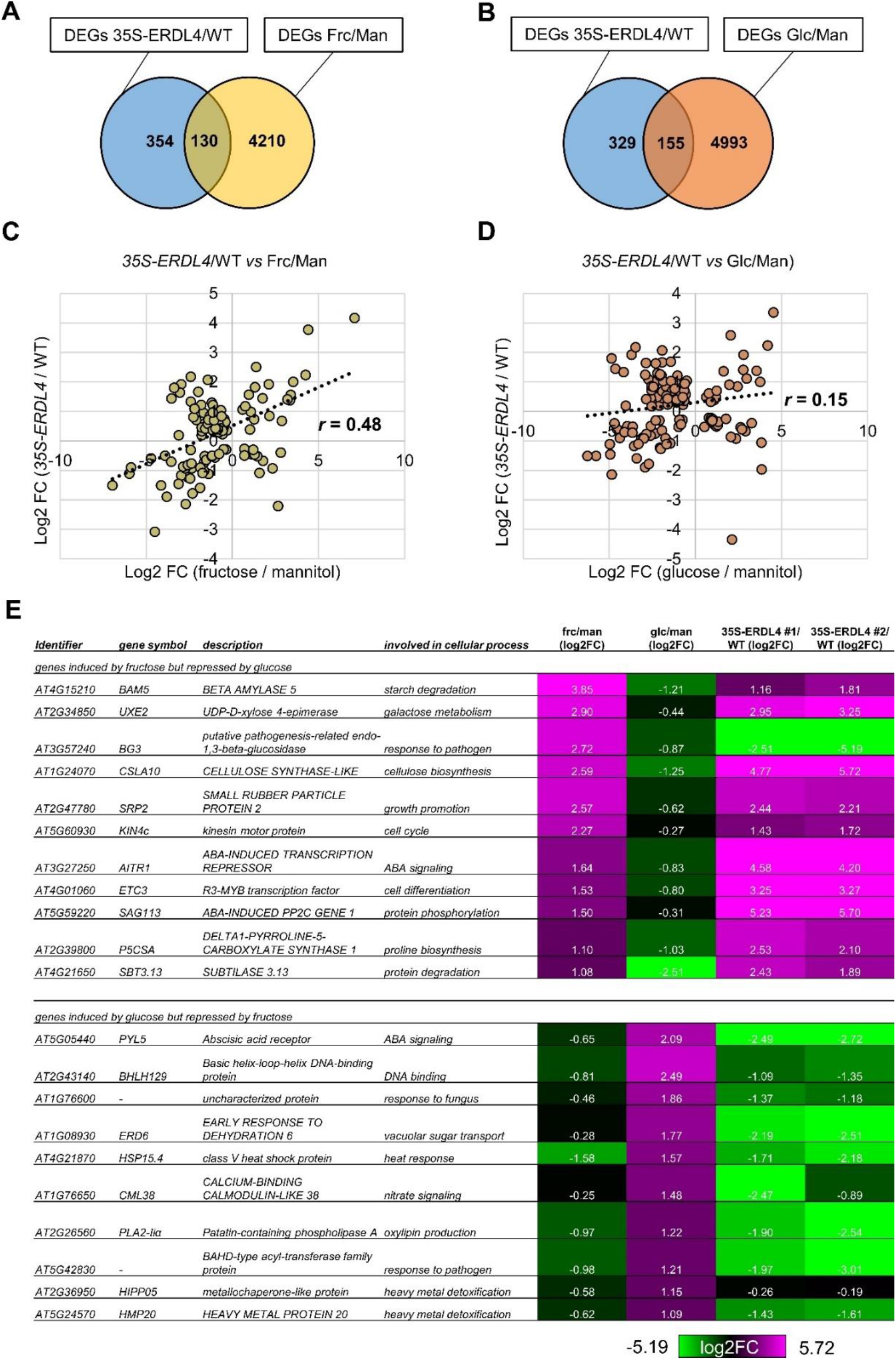
Venn-diagram and correlation analysis between differentially expressed genes (DEGs) from *ERDL4-* overexpressing and WT plants and from fructose (Frc) or glucose (Glc) and mannitol (Man = control) incubated leaf discs. **A)** and **B)** show Venn-diagrams of DEGs. Numbers within circles represent the number of DEGs (significant at p < 0.001 according to the double-sided Students *t*-test of three replicates). There were 484 *35S-ERDL4*-dependent DEGs, 4340 Frc-dependent DEGs, and 5148 Glc-dependent DEGs. **A)** Diagram showing overlap of DEGs from *35S-ERDL4* and Frc-treatment. There were 130 DEGs regulated by both *ERDL4*-overepression and Frc. **B)** Diagram showing overlap of DEGs from *35S-ERDL4* and Glc-treatment. There were 155 DEGs regulated by both *ERDL4*-overexpression and Glc. **C)** Correlation between relative expression (based on log2-fold-change) of 130 DEGs between 35S-ERDL4 (against WT) and Frc (against Man). *r* = Pearson coefficient calculated from ERDL4/WT and Frc/Man matrices **D)** Correlation between relative expression (based on log2-fold-change) of 155 DEGs between 35S-ERDL4 (against WT) and Glc (against Man). *r* = Pearson coefficient calculated from ERDL4/WT and Glc/Man matrices. **E)** List of DEGs oppositely regulated by frc or glc ranked after their Log2FCs. The upper part of the table lists the 10 most strongly frc-induced genes that were at the same time repressed by glc and the lower part the 10 most strongly glc-induced genes that were at the same time repressed by frc (Log2FCs cutoff ≤ 0.25).

Next, we extracted two subsets of DEGs from the sugar feeding data set: (i) the ten strongest fructose-upregulated DEGs that were at the same time repressed by glucose (cut off: log2FC ≤ −0.25) and (ii) the ten strongest glucose-upregulated DEGs that were at the same time repressed by fructose (cut off: log2FC ≤ −0.25). These genes served as indicators for cytosolic fructose or glucose accumulation, respectively. Since the NAF data (**Figure 5**) indicated that both fructose and glucose levels increased in the cytosol of *35S-ERDL4* plants, the expression of these genes may provide information about the effectiveness of both sugars evoking a directed response on gene expression when present in elevated levels at the same time.

Nine of the ten genes which were fructose-induced but glucose-repressed were also clearly induced in *35S-ERDL4* lines (with log2FCs > 1) except for *A73G57240*, a putative pathogenesis-related endo-1,3-beta-glucosidase-encoding gene which was markedly downregulated in both *ERDL4* overexpressing lines (**Figure 6E**). Conversely, all ten DEGs that were increased by glucose but repressed by fructose showed also clear downregulation in the *35S-ERDL4* lines (**Figure 6E**). These results showed that the elevated cytosolic fructose levels in *35S-ERDL4* plants – although present at a lower overall concentration than that of glucose – had a stronger regulatory effect on the expression of these markers than the simultaneously increased cytosolic glucose and suggested that fructose constitutes a potent signal on gene expression which could in certain cases override an oppositely directed glucose signal.

### High sugar content of ERDL4-overexpressors depends on TST proteins

The previous results suggested that ERDL4 acted as a vacuolar sugar transporter catalyzing release of fructose from vacuoles into the cytosol and triggering fructose-specific gene expression responses. If ERDL4 would exhibit such function, how could *35S-ERDL4* plants then achieve the observed strong accumulation of glucose in the vacuole? To address this question and because cytosolic sugar levels are connected to the expression of sugar-related genes (Strand *et al*., 1997; Smeekens, 1998; Pommerrenig *et al*., 2018; Chen *et al*., 2019; Vu *et al*., 2020), we checked the expression of the two major vacuolar sugar importer genes, *TST1* and *TST2* and that of *ERDL6* encoding a glucose/proton symporter of the tonoplast, in WT, *ERDL4*-overexpressing lines, and *erdl4*-knockout mutants (**Figure 7A–C)**. Of the two *TST* genes *TST2* was significantly upregulated in *ERDL4*-overexpressing but not *erdl4* knock out plants in comparison to the WT (**Figure 7A, B**). The expression of *ERDL6* on the other hand was unchanged in the *35S-ERDL4 lines*, but greatly increased in *erdl4-1* and *erdl4-2* knockouts (**Figure 7C**). These results suggested that the observed increased glucose content of vacuoles from *35S-ERDL4* plants was a result from TST-derived sugar/proton antiport into vacuoles which did not occur in the same manner in *erdl4* knockouts where *TST* expression was unaltered or even reduced.

**Figure 7.**
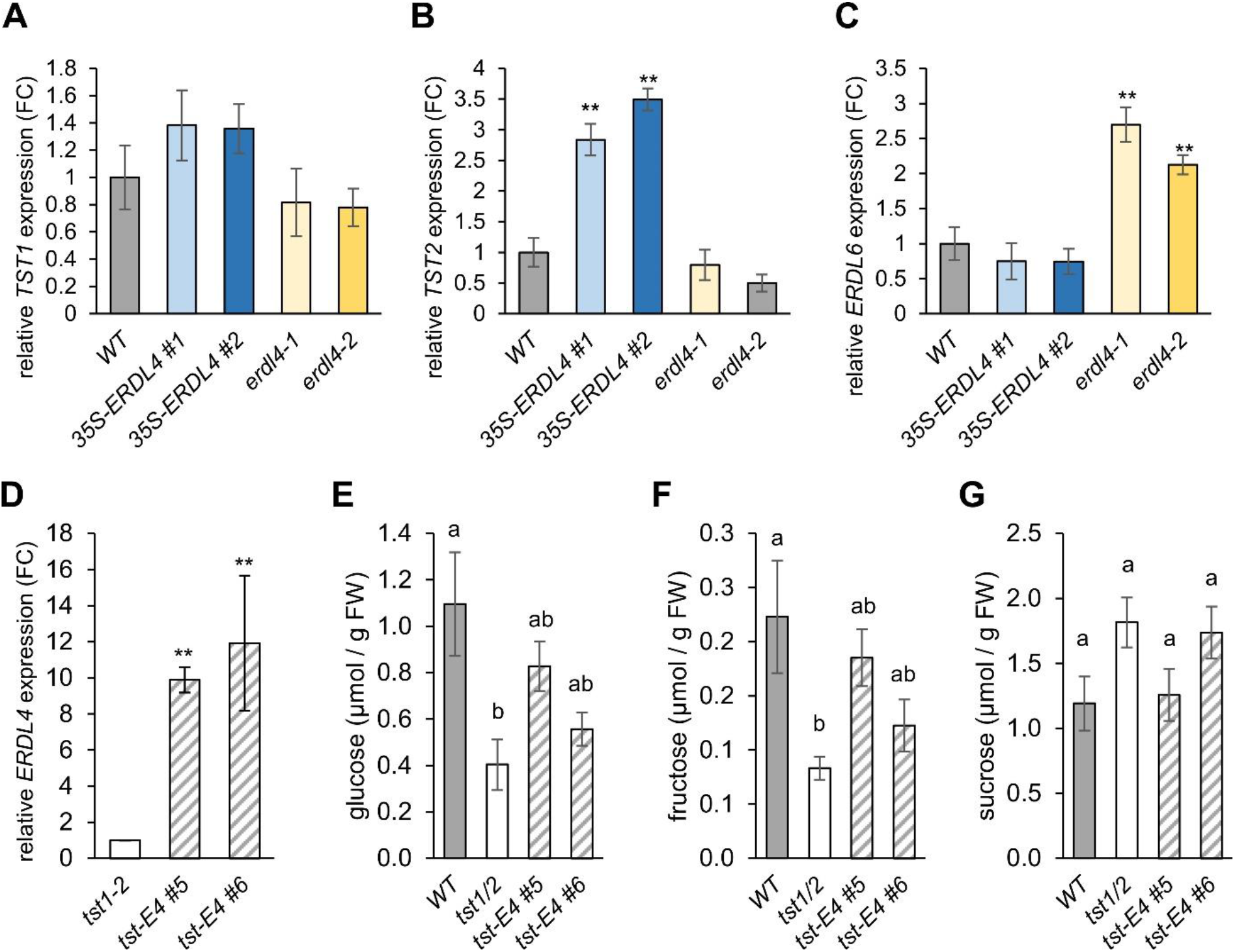
Expression of vacuolar sugar transporters in plants with altered *ERDL4* expression and overexpression of *ERDL4* in *tst1/2* double knockouts. **A-C)** Expression of *TST1* (A), *TST2* (B), *ERDL6* (C) in *ERDL4*-overexpresing and erdl4 mutant plants. **D)** Relative expression of *ERDL4* in *tst1/2* and *tst-ERDL4* plants determined by RT-qPCR. Bars show means from n=4 homozygous plants ± SE. Asterisks indicate significant changes according to Students t-test in comparison to the control (*tst1/2*) (** p < 0.01, *** p < 0.001). **E-F)** Contents of glucose (B), fructose (C), sucrose (D) in shoots of WT, tst1/2 double knockout and two tst-ERDL4 lines. Bars are means from n=5 plants ± SE. Different letters above bars indicate significant differences according to One-Way ANOVA with post-hoc Tukey HSD testing (p < 0.05).

To confirm this hypothesis, we expressed the *35S-ERDL4* construct in *tst1/2* double knock-out plants (*tst-ERDL4 #5* and *tst-ERDL4 #6*). The *tst1/2* mutant lacks the two major proteins transporting sugar from the cytosol into the vacuolar lumen, TST1 and TST2 (Wormit *et al*., 2006; Schulz *et al*., 2011; Vu *et al*., 2020). The expression of *ERDL4* in *tst-ERDL4 #5* and *#6* plants exceeded that of *tst1/2* control plants ten- and twelvefold, respectively (**Figure 7D**). The *tst1/2* double knockout plants had significantly lower glucose and fructose levels (and slightly increased levels of sucrose) in comparison to WT plants (**Figure 7E–G**), confirming results from previous analyses (Klemens et al., 2014; Vu et al., 2020). When *ERDL4* was overexpressed in these double mutants (as in *tst-ERDL4* #5 and *tst-ERDL4 #6*), glucose and fructose levels of such plants did not exceed those of wild type plants (**Figure 7E, F**). This result showed that ERDL4 was not directly involved in the transport of sugars into the vacuole and strongly indicated that TST1 and TST2 were the major factors responsible for the massive monosaccharide accumulation in the *35S-ERDL4* lines (**Figure 7A**). However, *tst-ERDL4* plants accumulated slightly higher glucose and fructose levels than *tst1/2* plants, suggesting that factors other than TST1 and TST2 contributed fractionally to vacuolar monosaccharide accumulation in *ERDL4* overexpressing plants.

### Fructose regulates key factors of vacuolar sugar homeostasis

Activity of TSTs have been shown to be regulated by their phosphorylation patterns and activity of specific kinases like VIK (for TST1; Wingenter *et al*., 2011) or CBL-INTERACTING PROTEIN KINASE 6 (CIPK6) (for TST2; Deng *et al*., 2020). CALCINEURIN B-LIKE PROTEIN 2 (CBL2)- dependent interaction of CIPK6 with TST2 has been shown to regulate sugar homeostasis in cotton (*Gossypium hirsutum*) (Deng *et al*., 2020). Expression of the TST1-kinase gene *VIK* was unaltered in WT and *ERDL4*-overexpressing plants (**Figure 8A**).

**Figure 8.**
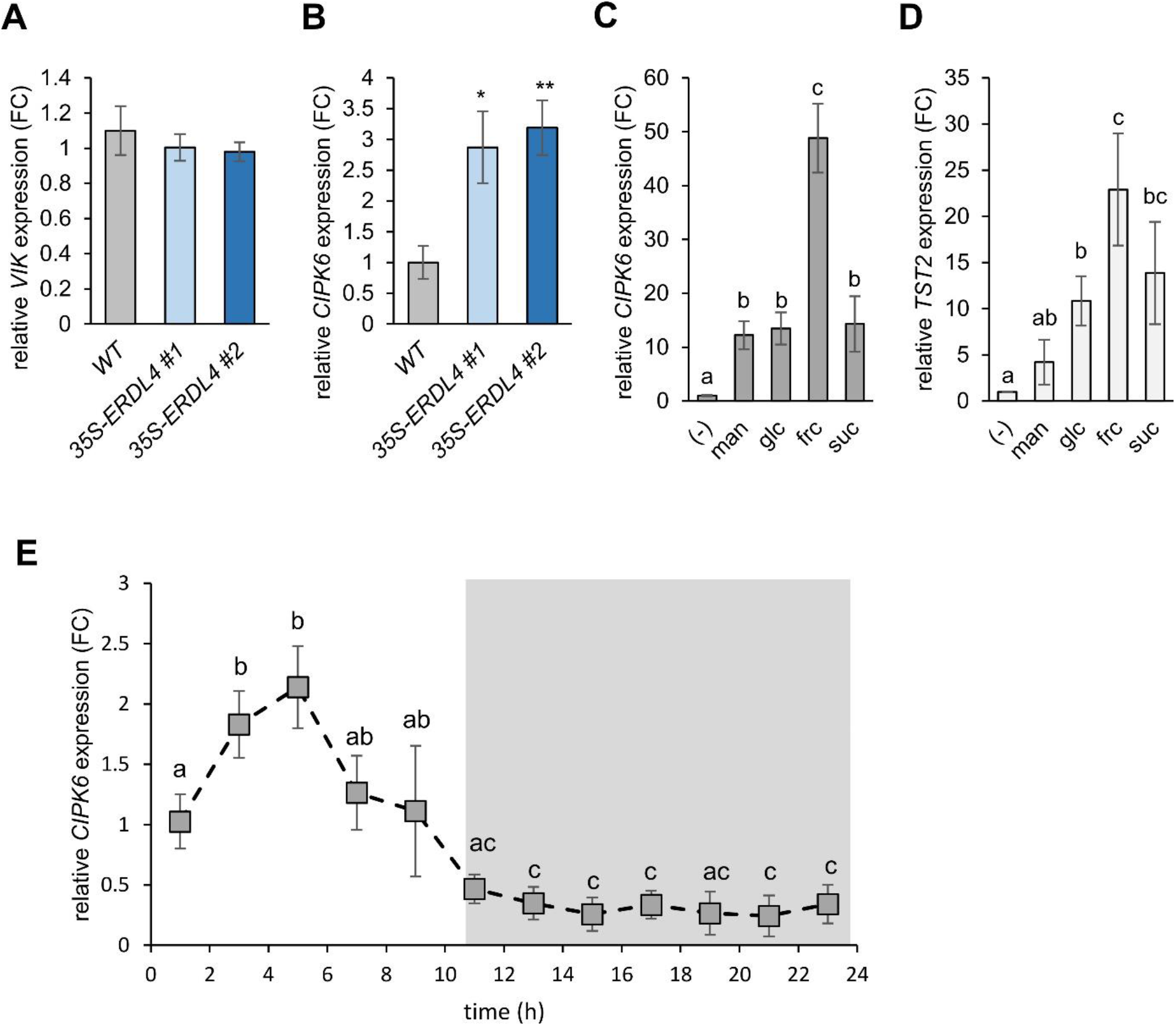
Sugar- and *ERDL4*-dependent expression of *VIK, CIPK6* and *TST2* based on RT-qPCR assays. **A)** relative expression of *VIK* in WT and *35S-ERDL4* plants. **B)** relative expression of *CIPK6* in WT and *35S-ERDL4* plants. **C, D)** relative expression of *CIPK6* **(C)** or *TST2* **(D)** in leaf discs incubated overnight in liquid MS medium supplemented without (-) or with 1% of the indicated sugars or mannitol (man). **E)** diurnal expression of *CIPK6*. Grey shading indicates dark phase (night time)

Phylogenetic analysis using sequences from the 26 *At*CIPKs and *Gh*CIPK6 identified *At*CIPK6 as closest homolog to *Gh*CIPK6 (**Supplemental Figure S4A**). Fructose accumulation has been associated with increased cytosolic Ca^2+^ levels through activation of voltage gated Ca^2+^ channels at the plasma membrane (Furuichi *et al*., 2001). The rise of Ca^2+^ levels might initiate downstream Ca^2+^ signaling leading to expression of genes responsible for sugar homeostasis like the CBL-CIPK pathway. Log2-fold changes from our RNA-Seq data indicated that six *CIPK* genes were highly regulated in presence of glucose and fructose relative to the control (**Supplemental Figure S4B**). Two of them, *CIPK9* and *CIPK20* were downregulated by both glucose and fructose and were similarly regulated in *35S-ERDL4* lines (**Supplemental Figure S4B**). In contrast, *CIPK7* and *CIPK25* were upregulated by the two sugars as well as in *35S-ERDL4 #1* and *35S-ERDL4 #2* (**Supplemental Figure S4B**). Only *CIPK5* and *CIPK6* showed fructose-specific upregulation and were also induced in both *ERDL4*-overexpression lines (**Figure 8B and Supplemental Figure S4B**). Although exhibiting strong relative induction *CIPK5* was expressed to a markedly lower extent when compared to *CIPK6* which makes CIPK6 a more likely target, responsible for TST2 phosphorylation and activation (**Supplemental Figure S4**). RT-qPCR confirmed the elevated *CIPK6* expression in *35S-ERDL4* (**Figure 8B**) as well as the induction of CIPK6 and TST2 by fructose (**Figure 8C, D**). When we checked the diurnal expression of *CIPK6* we found that *CIPK6* expression patterns closely overlapped with that of *TST2* (**Figure 2 and Figure 8E**) supporting a possible interaction of CIPK6 and TST in Arabidopsis during the light phase.

### ERDL4 influences cold acclimation and freezing tolerance

Upon exposure to low temperatures, plants increase their overall sugar content in leaves drastically within a comparably short time, usually within one to three days, in a process known as cold acclimation (Klemens *et al*., 2013; Rodrigues *et al*., 2020; Ho *et al*., 2020). In Arabidopsis, cold acclimation-dependent sugar accumulation occurs mainly in vacuoles and depends on TST proteins (Klemens et al., 2013, Vu et al., 2020).

When we checked for expression of *ERDL4* in cold-acclimated WT plants (three days at 4°C) we found that *ERDL4* mRNA increased almost 20-fold in comparison to plants grown in control (20°C) conditions (**Figure 9A**). This suggested that vacuoles of cold-acclimated plants released fructose to a greater extent than vacuoles of plants grown under control conditions (20°C) mimicking a situation existing in *35S-ERDL4* plants (**Figure 5**). Because *ERDL4-*overexpressing plants and *erdl4* knockout mutants accumulated higher and lower levels of monosaccharides, respectively, in comparison to wild types already at 20°C (**Figure 5A**), we were interested to check sugar accumulation in these lines at 4°C (**Figure 9**). After 48h in the cold, all lines exhibited high glucose, fructose, and sucrose levels in leaves (**Figure 9B–D**) exceeding those of 20°C-grown plants markedly (**Figure 5A**). Glucose accumulated to similar amounts in WT, *35S-ERDL4 #1* and *#2* (between 20.3 and 21.9 μmol/g FW) (**Figure 9B**). Plants from *erdl4-1* and *erdl4-2* knockout lines accumulated only about 60% (12 μmol/g FW) and 50% (10.1 μmol/g FW) in comparison to wild-types, respectively (**Figure 9B**). Fructose and sucrose accumulation was not altered significantly between the tested lines (**Figure 9C, D**).

**Figure 9.**
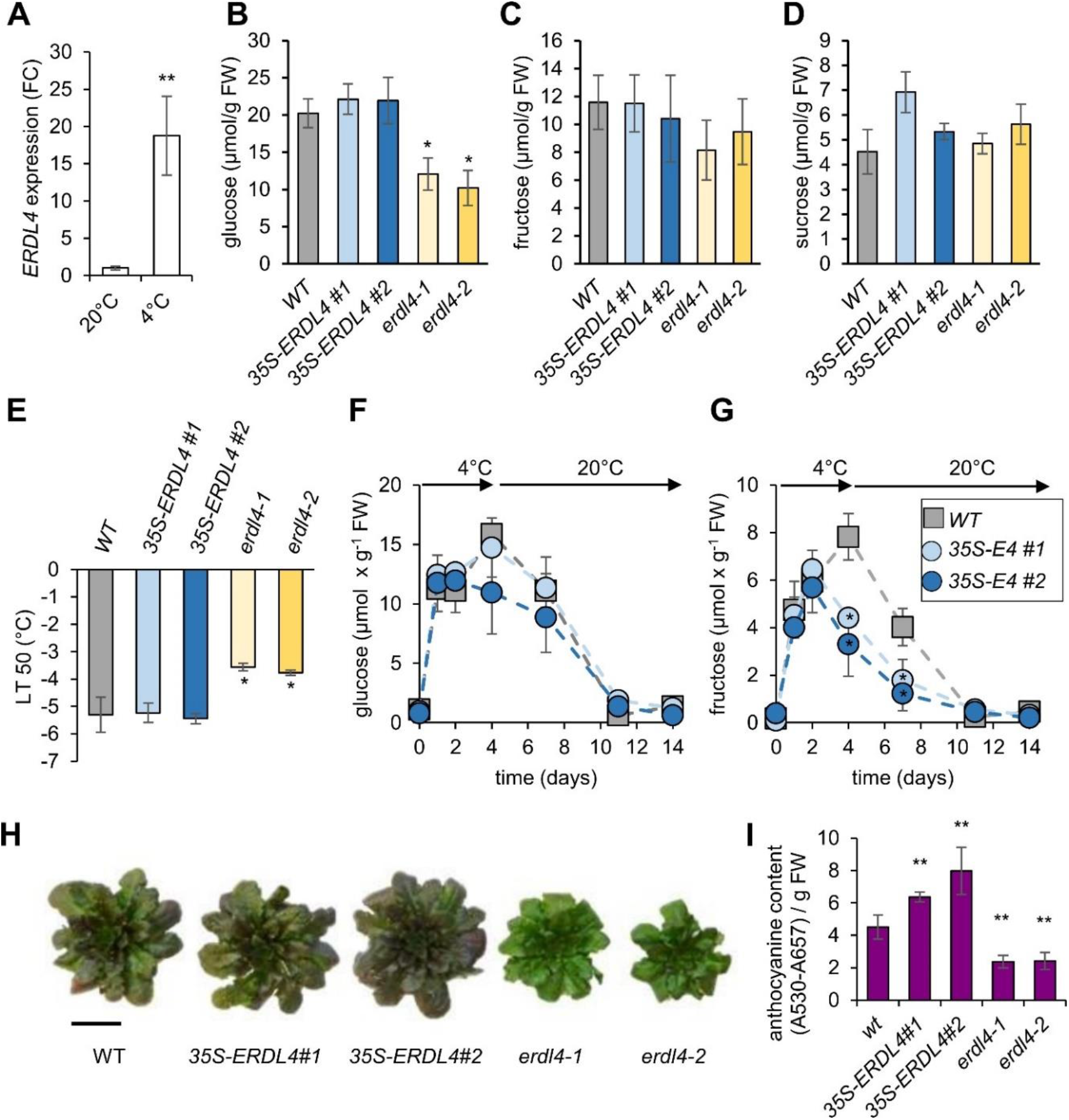
Cold treatments of *ERDL4*-overexpressing and *erdl4* knock-out plants. **A)** Expression of *ERDL4* relative to *ACT2* in WT plants grown under control (20°C) or for three days at 4°C. Bars are means from n=4 individual plants ± SE. Asterisks indicate significant differences according to Students t-test (** = p < 0.005) **B-D)** contents of glucose **(B)**, fructose **(C)** and sucrose **(D)** in rosette leaves of wild-type (WT), *ERDL4*-overexpressing (*35S-ERDL4 #1, #2*) and knockout (*erdl4-1, erdl4-2*) plants grown for 48h at 4°C. Asterisks indicate significant differences to WT according to Students t-test (p < 0.05) **E)** electrolyte leakage assay with leaf samples taken from plants grown for three weeks at 20°C and then four days at 4°C. Asterisks indicate significant differences to WT (p < 0.05). **F, G)** glucose (F) and fructose (G) accumulation of WT and *35S-ERDL4* plants during cold acclimation (until day 4) and de-acclimation (until day 14). **H, I)** long-term response to low temperatures. WT, *35S-ERDL4* and *erdl4* knockout plants were grown at 4°C for 14 days and harvested for determination of anthocyanin content **H)** exemplary pictures of rosettes. Bar = 2 cm. **I)** anthocyanin content of leaves. Bars represent mean values from n=4 plants ± SE. Asterisks indicate significant differences to WT according to double-sided Students t-test (** = p < 0.005).

The differential sugar accumulation of the analyzed lines in the cold had consequences for their freezing tolerance as revealed by electrolyte leakage assay conducted with leaf samples of plants grown for three weeks at 20°C and then 48h at 4°C. Leaves from such cold-acclimated plants were cooled to - 10°C in water. After thawing on ice overnight, electric conductivity serving as an indicator for the number of ions released from ruptured cells was measured (Sukumaran and Weiser 1972). The temperature at which 50% of the ions were released (LT50) of *erdl4-1 (*−3.5°C) and *erdl4-2* (−3.7°C) knockout plants was significantly higher than that of WT (−5.3°C) and *35S-ERDL4* (−5.2 and −5.4°C) plants (**Figure 9E**). The data indicated that *erdl4* mutants were less freezing-tolerant than WT and *35S-ERDL4* plants and suggested that the lower overall sugar accumulation in *erdl4* mutants may be causative for the higher freezing sensitivity. Interestingly, *35S-ERDL4* plants exhibited different fructose accumulation kinetics during de-acclimation, that is when plants were transferred from 4°C acclimation phase back to 20°C (**Figure 9F, G**). During such phase, plants remobilized and utilized vacuolar-stored glucose and fructose to reach pre-acclimation levels within seven to ten days (**Figure 9F, G**). Decrease of fructose during de-acclimation occurred faster in *35S-ERDL4* plants when compared to WT resulting in altered fructose to glucose ratios: While WT_frc/glc_ ratios at day four of acclimation and day three of de-acclimation were about 0.52 and 0.36, respectively, *35S-ERDL4 #1*_frc/glc_ (0.29 and 0.15) and *35S-ERDL4 #1*_frc/glc_ (0.30 and 0.13) ratios at these time points were markedly lower. The shifted ratios in *ERDL4*-overexpressing lines in favor of glucose supported a scenario where fructose was increasingly released from vacuoles by increased ERDL4 activity during the late acclimation and de-acclimation phases.

In contrast to short-time acclimation responses, plants transit into the so-called adaptation phase after prolonged exposure to environmental stress (Stitt and Hurry, 2002). Adaptation to cold is associated with accumulation of anthocyanins acting as protectants of cellular compounds against oxidative stress. After 14 days of cold treatment, wild-type and *35S-ERDL4*, but not *erdl4* knockout plants exhibited purple coloration of their leaves indicating accumulation of anthocyanins. Moreover, *erdl4* knockout plants exhibited markedly reduced rosette diameter in comparison to WT and *35S-ERDL4* plants at this stage (**Figure 9H**). The anthocyanin content of *35S-ERDL4* lines exceeded that of wild-type plants by 40% (*35S-ERDL4 #1*) and 75% (*35S-ERDL4 #2*). The anthocyanin contents of *erdl4* knockouts on the other hand reached only about half of the wild-type values (**Figure 9I**). The data suggested reduced efflux of monosaccharides from vacuoles of *erdl4* plants and indicated that sugars released from vacuoles during cold adaptation were used as building blocks for different anthocyanidins serving as factors for maintaining growth during low temperatures.

### ERDL4 influences flowering, silique development, seed size, and seed lipid content

One of the most interesting plant properties is the final yield of fruits and storage organs. Thus, to get first insight into putative effects of ERDL4 activity on yield, we checked the effect of differential *ERDL4* expression on Arabidopsis inflorescence and seed characteristics (**Figure 10**). It turned out that *35S-ERDL4* plants flowered later than WT plants but grew longer inflorescences with more branches (**Figure 10A, B**). Altered *ERDL4* expression also influenced silique and seed properties. Siliques of *35S-ERDL4* plants were slightly longer in comparison to those of wild type plants (**Figure 10C**). The number of seeds of such longer siliques of *35S-ERDL4* plants was not different from that of WT siliques (**Figure 10D**). However, seeds of *35S-ERDL4* plants were bigger than wild-type (**Figure 10F**) and had a 20% (*35S-ERDL4 #1*) to 30% (*35S-ERDL4 #2*) higher 1000-seed weight when compared to wild-type (**Figure 10E**). Siliques of both *erdl4* mutants on the other hand were shorter (**Figure 10C**) and contained significantly fewer seeds than wild type **(Figure 10D**). Both *erdl4* mutants produced wrinkled seeds (**Figure 10F**) with significantly diminished 1000-seed weights (**Figure 10E**). The altered seed weight and shape was also reflected by different seed lipid contents in *35S-ERDL4* plants and *erdl4* mutants. While seeds of *35S-ERDL4* lines had significantly higher lipid content, *erdl4* mutants showed reduced contents when compared to WT seeds (**Figure 10G**). These results showed that ERDL4 was important for proper seed production and suggested that intracellular sugar compartmentation influenced both silique growth and seed filling.

**Figure 10.**
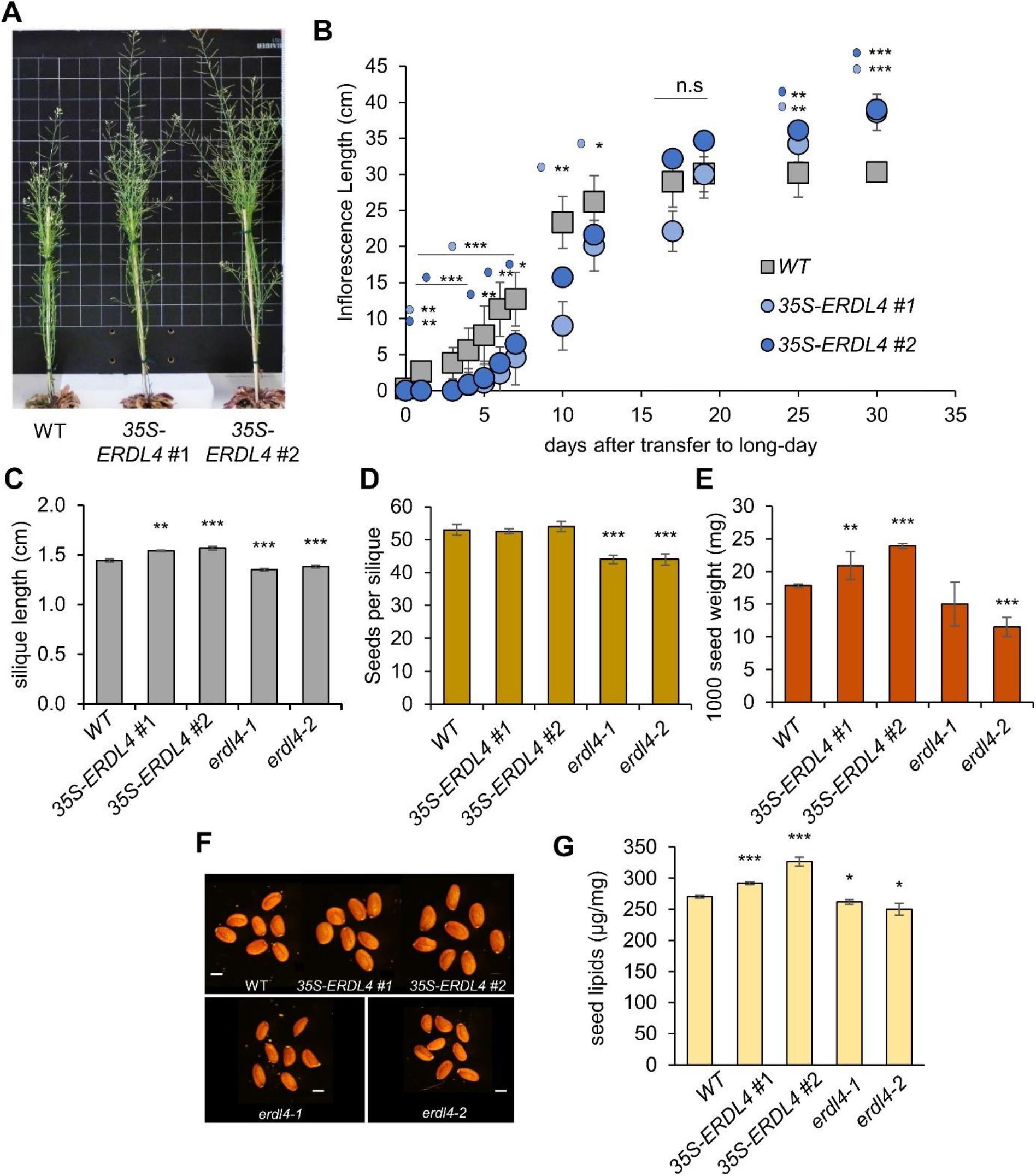
Phenotypes of *ERDL4* mutants during the generative phase. **A)** Exemplary phenotypes of *ERDL4-* overexpressing plants during flowering. **B)** Recording of inflorescence length of WT and two *ERDL4*-overexpressing lines after transfer from short-day to long-day conditions. Each data point represents the average main inflorescence stem length of at least 20 plants ± SE. **C)** Silique length of WT and two *ERDL4*-overexpressing lines. Bars represent means from 30 siliques from ≥15 plants ± SE. **D)** Number of seeds per silique of WT and two ERDL4-overexpressing lines. Bars represent means from 20 siliques from 10 plants ± SE. **E)** Thousand-Seed weight of dry mature seeds from WT, *erdl4* knockout mutants and *ERDL4-*overexpessing plants. Bars represent means from 1000-Seed weights of n=6 plants per line. Asterisks indicate significant differences according to Students t-test (** = p<0.01; *** = p < 0.001). **F)** Mature seed phenotypes from WT, *ERDL4*-overexpressing, and *erdl4* knock-out mutant plants. Representative pictures are shown. **G)** Seed lipid content of dry mature seeds from *erdl4*-knock-out mutants and *ERDL4-*overexpessing plants. Bars represent means from n=6 plants per line. Asterisks indicate significant differences according to Students t-test (* = p < 0.05; ** = p<0.01; *** = p < 0.001).

## Discussion

Sugars provide cellular energy and carbon precursors for most anabolic reactions. In addition, sugars affect early plant germination, development of the vasculature or govern onset of flowering. It is well known that sugars are key components controlling yield of fruits and storage organs and influence positively plant tolerance to abiotic stresses like drought or cold temperatures (Ruan, 2014; Le Hir *et al*., 2015; Julius *et al*., 2017; Pommerrenig et al., 2020; Keller *et al*., 2021).

For most of these functions plant cells must monitor the levels of sugars. The molecular sensor leading to glucose perception is well understood since this sugar is recognized via its interaction with cytosolic HEXOKINASE1 (HXK1) (Jang *et al*., 1997; Rolland *et al*., 2006). The subsequent signal transduction pathway comprises altered phosphorylation of selected transcription factors (Hu *et al*., 2016) and nuclear interaction of HXK1 with partner proteins (Cho *et al*., 2006). In case of fructose, effects of this sugar on e.g., root morphology (Zhong *et al*., 2020; Valifard *et al*., 2021) or plant-pathogen interaction (Lecompte *et al*., 2017) as well as fructose transport activity of plasma membrane-(Nørholm *et al*., 2006; Rottmann *et al*., 2018) or tonoplast-located proteins (Wormit *et al*., 2006; Chardon *et al*., 2013; Guo *et al*., 2014) have been reported. However, because glucose, fructose, and sucrose metabolism are in most cases linked via the action of sucrose biosynthesis and degrading enzymes (Volkert *et al*., 2014; Vu *et al*., 2021), data on fructose-specific gene expression and specific fructose sensing mechanisms are scarce. In fact, for fructose signaling a sensor protein is still lacking and only two proteins are proposed to be intermediates of the fructose signaling pathway, namely the transcription factor ANAC89 and a fructose1.6-bisphosphate phosphatase, as revealed on the mutant FRUCTOSE INSENSITIVE1 (FINS1) (Cho and Yoo, 2011; Li *et al*., 2011). Unfortunately, ANAC89 function seems exclusively limited to the Cape Verde Islands (Cvi) Arabidopsis population (Li *et al*., 2011) and for neither ANAC89 nor FINS1 fructose binding has been shown so far.

Given the fact that total fructose levels of plant leaf tissue are generally five to ten-times lower than that of glucose (Schofield *et al*., 2009; Patzke *et al*., 2019; Pommerrenig *et al*., 2019; Vu *et al*., 2020; and **Figure 5**), it is reasonable to assume a stringent regulation of free fructose pools in plant cells. This assumption is supported by the negative effects of imbalanced subcellular fructose homeostasis, as present in knock-out mutants of the vacuolar fructose transporter SWEET17, on stem morphology, root architecture and abiotic stress tolerance (Aubry *et al*., 2021; Valifard *et al*., 2021). A specific fructose function is also indicated by the observation that this sugar can induce expression of genes encoding transcription factors involved in lateral root development inducing corresponding morphological changes and increased drought tolerance (Valifard *et al*., 2021).

Clearly, plants need therefore to be able to regulate cytosolic fructose levels to adjust gene expression and corresponding plant responses. In addition to the cytosol, glucose and fructose can accumulate in vacuoles as sucrose hydrolysis products (Vu *et al*., 2020). Accordingly, by action of either glucose- or fructose-specific vacuolar transporters, individual sugars can be sequestered into or released from the vacuole in a differential and coordinated manner to evoke specific molecular and physiological responses.

Our observations that the vacuolar transporter ERDL4 is involved in fructose transport and fructose-related responses is fully in line with the hypothesis that plant cells adjust their sugar homeostasis by regulating cytosolic fructose levels via vacuolar transporters. We provide evidence that changes of cytosolic fructose levels as provoked by either overexpression or knockout of *ERDL4* steered fructose-dependent changes of gene expression resulting in altered source and sink growth, accompanied by changed sugar accumulation and transport in leaves.

ERDLs constitute the largest MST subfamily and until now only a few members have been functionally characterized. *ERD6* from Arabidopsis, the first *ERDL* gene to be identified, has been demonstrated to be induced by dehydration and cold stress (Kiyosue *et al*., 1998). GFP-fusions of the yet characterized ERDLs all localize to the tonoplast (Yamada *et al*., 2010; Poschet *et al*., 2011; Remy *et al*., 2014). The ERDL4 protein analyzed in this work has been detected together with its homologs ERDL7 and ERDL8 in tonoplast membrane fractions by LC-MS/MS-based proteomics analysis (Pertl-Obermeyer *et al*., 2016). Analysis of the localization of an ERDL4-GFP fusion in mesophyll protoplasts clearly indicated that ERDL4 localizes to the tonoplast corroborating these findings (**Figure 1)**. Given the overlapping subcellular localization of ERDLs and tissue- and stress-dependent expressing patterns of the *ERDL* genes (Slawinski *et al*., 2021b), it is tempting to speculate that individual ERDLs differ in their substrate specificities and affinities. Indeed, ERD6 and its close homolog ERD6-like 3 (ERDL3) have been attributed as low-affinity, broad-spectrum facilitators for glucose and other monosaccharides like fructose, xylose, mannose, and galactose (Yamada *et al*., 2010). ERDL7 mediated zinc efflux when expressed in yeast cells (Remy *et al*., 2014) and the closest homologs to ERDL4, ERDL6 from Arabidopsis and *Bv*IMP from sugar beet, have been characterized as proton-coupled glucose transporters (Poschet *et al*., 2011; Klemens *et al*., 2014). Our conclusion that ERDL4 preferentially mediated import of fructose from the vacuolar lumen into the cytosol resulted from different lines of evidence: Firstly, uptake experiments on *ERDL4*-expressing *Xenopus laevis* oocytes suggested that ERDL4 transports fructose but not glucose or sucrose (**Figure 1**). Secondly, subcellular distribution of glucose and fructose in *35S-ERDL4* differed from that in WT plants exhibiting significantly elevated levels in the cytosol of overexpressors in comparison to WT (**Figure 5**). Thirdly, the increased levels of glucose and fructose in the cytosol of *35S-ERDL4* plants stimulated expression of specific glucose- and fructose-responsive genes substantiating results of the NAF analysis (**Figure 5, 8**).

The recording of sugar-responsive gene expression patterns by RNASeq allowed for the identification of genes which were oppositely regulated by glucose and fructose (**Figure 6**). Based on the combined analysis of *ERDL4*-overexpressing mutants and the comparative analysis of gene expression with feeding of various sugars **(Figure 6),** we raised substantial evidence that ERDL4 stimulated fructose-specific signaling processes which govern development and size of various Arabidopsis organs including seeds and roots. The altered subcellular monosaccharide levels and compartmentation of *ERDL4*-overexpressing lines lead to morphological, physiological, and molecular alterations of these plants in comparison to wild-types (**Figure 3, 9, 10**). Surprisingly, we found that plants of these lines were larger and accumulated sugar to higher amounts when compared to corresponding WT plants. In this regard, they resembled *TST*-overexpressing (Wingenter *et al*., 2011) and *MdERDL6-overexpressing* (Zhu *et al*., 2021), rather than *ERDL6*-overexpressing plants (Poschet *et al*., 2011). Latter plants which overexpressed the close glucose/proton symporting homolog to ERDL4 exhibited high cytosolic glucose levels, reduced shoot biomass, and impaired freezing tolerance (Poschet *et al*., 2011; Klemens *et al*., 2014).

Modulation of sugar homeostasis by altering sugar transporter levels leads to differential regulation of other sugar transporters probably for maintaining internal sugar balance (Sun *et al*., 2019; Zhu *et al*., 2021). Interestingly, a similar regulation is also observed in *ERDL4*-overexpressing plants where high cytosolic fructose and glucose levels result in differential regulation of the vacuolar transporter gene *TST2* (**Figure 7**). However, the strong fructose induction of *TST2* expression (Wormit *et al*., 2006 and **Figure 8**) and the fact that *ERDL4-*overexpression in *tst1/2* knockout background did not result in high leaf sugars (**Figure 7**) allows to conclude that the increased vacuolar glucose accumulation in *ERDL4-*overexpressing lines (**Figure 5**) was mainly due to the increased *TST2* levels.

Recent analysis of *ERDL* genes revealed their early emergence and conservation within the plant kingdom (Slawinski *et al*., 2021a). The *ERDL4* and *ERDL6* genes are expressed in almost all organs (Slawinski *et al*., 2021b). However, as we show here, the two genes have non-overlapping/opposite diurnal expression patterns **(Figure 2)**. Expression of *ERDL4* increases at the onset of night and reaches its maximum at dawn, whereas *ERDL6* levels are highest at midday and in the evening (**Figure 2**) suggesting different functions of the two proteins. Diurnal expression of *ERDL4* concurs with a vacuolar export function essential for mobilization of vacuole-stored sugars in the dark (Stitt *et al*., 1985; Schulz *et al*., 2011; Vu *et al*., 2020). The observation that expression of vacuolar sugar importers, *TST1* and *TST2*, rise during the light phase and decline in the dark supports the notion that during the light phase, sugars are accumulated in the vacuole for which induction of sugar importers is essential (**Figure 2**).

Vacuolar sugar transport can additionally be regulated by phosphorylation of TST proteins via either the Arabidopsis protein kinase VIK (Wingenter *et al*., 2011) or the cotton protein Calcineurin B-like protein (CBL) interacting protein kinase 6 (CIPK6) (Deng *et al*., 2020). In cotton TST2, Ser^448^ has been found as a target amino acid being phosphorylated by *Gh*CIPK6 (Deng *et al*., 2020). (Interestingly,) this residue is conserved in barley, rice, and Arabidopsis TST2 proteins (Hunter, 2020, Deng *et al*., 2020, Dhungana and Braun, 2021) suggesting conservation of this Ca^+^ and phosphorylation signalling resulting in sugar accumulation across these species. Interestingly, like *TST2, CIPK6* was strongly induced by fructose and to a weaker extent by glucose or mannitol (**Figure 8**). Like TSTs, CIPKs are centrally involved in the responses to several types of stress, including cold and drought (Pandey *et al*., 2015; Chen *et al*., 2021; Ma *et al*., 2021). However, future analyses of TST2 activity and sugar accumulation patterns of *cipk6* mutants as well as interaction studies of CIPK6 and TST2 are necessary to investigate a direct effect of CIPK6 on TST2 phosphorylation and activity.

*ERDL4* was also upregulated during cold stress (**Figure 9**). Arabidopsis plants are known to accumulate sugars during acclimation process which increases their frost tolerance (Wormit *et al*., 2006; Klemens *et al*., 2013). Monosaccharide accumulation in vacuoles in the cold occurs almost exclusively via monosaccharide and sucrose import by TST proteins and successive hydrolysis of sucrose by vacuolar invertases (Schulz *et al*., 2011; Klemens *et al*., 2013; Vu *et al*., 2020) resulting in a proportional increase of glucose and fructose moieties in the cold. Our observations support a scenario where cytosolic fructose acts as stimulant for vacuolar sugar accumulation and induction of both the major vacuolar sugar antiporter TST2 and its potential kinase CIPK6 (**Figure 8**). Consistent with a TST-activating function of ERDL4-mediated fructose release from vacuoles in the cold, *erdl4* knockout lines accumulated less monosaccharides in the cold, however their fructose to glucose ratio was roughly unaltered in comparison to the WT (**Figure 5 and Figure 9**). In *ERDL4-*overexpressing plants the fructose to glucose ratio was clearly lower, although these plants accumulated monosaccharides to similar levels as WT plants. The non-proportional decrease of glucose and fructose in *35S-ERDL4* plants in comparison to WT during de-acclimation suggested that fructose was released to a higher extent than glucose from vacuoles of *35S-ERDL4* plants and substantiated the assumption that ERDL4 acts as fructose rather than glucose transporter. These interpretations are fully in line with the strong increase of glucose but not of fructose in the vacuolar fraction of *ERDL4*-overexpressing plants in contrast to WT vacuoles (**Figure 5**). Under stress conditions, by such sugar species-selective release mechanism, fructose released from vacuoles to the cytosol may also provide carbon skeletons which can serve as a fuel in glycolysis and at the same time ensure vacuolar glucose accumulation.

In contrast to short time acclimation responses, plants transit to an adaptation phase after prolonged exposure to environmental stress. During long term cold adaptation, the metabolic processes in plants are different to acclimation. Here, plants are confronted with developing oxidative stress by reactive oxygen species build-up, possibly cell damaging processes, which are in part counter acted by accumulation of anthocyanins acting as protectants of cellular compounds (McKown *et al*., 1996). Whereas *35S-ERDL4* overexpressing lines accumulated anthocyanins to higher levels when compared to WT, *erdl4* knockout plants accumulated less anthocyanins. The observed effects in the cold suggest a fructose-dependent regulation of cold-induced accumulation of these protective compounds during short and long-term exposure to low temperatures and towards freezing.

In summary, our findings highlight the importance of cytosolic fructose levels to adjust plant growth and their tolerance to abiotic stress and shed light on fructose signaling and homeostasis in Arabidopsis, which surprisingly still represents a comparably “dark spot” in plant primary metabolism. Thus, it will be challenging to identify the corresponding sensor and signal transduction pathways translating dynamic changes of fructose levels into a coordinated cellular response.

## Materials and methods

### Plant material and growth conditions

Wild-type and mutant Arabidopsis (*Arabidopsis thaliana*) plants were grown in soil substrate (ED-73; Patzer; www.einheitserde.de) in a growth chamber (Weiss-Gallenkamp, Heidelberg, Gemany) at constant temperature and light of 22°C and 120 μmol quanta m^-2^ s^-1^ (μE) respectively. Plants were cultivated under short days (10 h light/14h dark regime. For cold treatment 6 weeks old plants grown under standard conditions were transferred to growth chamber at 4°C for 3 days. For root length analysis seeds were surface sterilized in 5% sodium hypochlorite and stratified for 24 h at 4°C before putting on ½ MS (pH 5.7) agar plates without sugars. Root to shoot ratios were determined from 6 weeks old plants grown in an already established hydroponic system (Conn et al., 2013) under standard conditions. For silique and inflorescence analysis plants were grown in standard conditions for 4 weeks and subsequently transferred to long day chamber (16 h light / 8 h dark).

### Cloning of *Arabidopsis ERDL4*-GFP construct

To determine the subcellular localization of ERDL4 in Arabidopsis, a *35S-ERDL4-GFP* fusion construct was generated using Gateway™ technology. The ERDL4-cds was amplified with Gateway-specific primers harboring the *attB1* and *attB2* sites ERDL4_gw_F (5’-ggg gac aag ttt gta caa aaa agc agg ctt aAT GAG TTT TAG GGA TGA TAA TAC G −3’) and ERDL4 _ gw_R (5’-ggg gac cac ttt gta caa gaa agc tgg gta TCT GAA CAA AGC TTG GAT C −3’). The resulting fragment was then introduced into Gateway entry vector pDON/Zeo and subsequently into destination vector pK7FWG2.0 (Karimi et al., 2002) via BP and LR reactions respectively. *Agrobacterium tumefaciens* strain GV3101 was used for the transformation of Arabidopsis plants via the floral dip method (Clough and Bent, 1998).

### Oocyte uptake experiments

For expression in oocytes, *ERDL4-cds* was cloned into the transcription vector pGEM-HE-Juel via BamHI and XbaI restriction enzyme sites. For cRNA synthesis, the construct was linearized with NotI. cRNA was synthesized using mMESSAGE mMACHINE T7 Ultra Kit (Carlsbad, CA 92008 USA).

The Xenopus laevis oocytes (defolliculated) were obtained from Ecocyte Bioscience (https://ecocyte-us.com/products/xenopus-oocyte-delivery-service/). The oocytes were kept in Oocyte Reagent2 (O-R2, 5 mM HEPES, 82.5 mM, NaCl, 2.5 mM KCl, 1 mM CaCl_2_, 1 mM MgCl_2_, 3.8 mM NaOH, and 1 mM Na2HPO_4_, pH 7.8) supplemented with Streptomycin (25 mg/L) and Gentamycin (25 mg/L) at 16°C. Healthy looking oocytes were injected with 50 ng of cRNA (determined by absorbance at 260 nm) in 50 nl water or with 50 nl water alone as a control using a glass capillary attached to a semi-automated micro-injector (World Precision Instruments, Inc. Sarasota, FL, USA) according to Aguero et al., 2018. The injected oocytes were incubated in O-R2 at 16°C for 3 days. Functional expression of the ERDL4 was assayed by the uptake of D-[^3^H(G)]-Fructose, D-[^14^C(U)]-Glucose and [^14^C(U)]-Sucrose (Hartmann Analytic, Braunschweig, Germany) for 30 min at 22°C in O-R2 buffer (pH 5.0) without antibiotics supplemented with 2 mM cold and 2.5 μCi radio-labelled sugar. For each condition, 10 oocytes were assayed. Oocytes were washed three times with 4 mL of ice-cold un-labelled O-R2 buffer. Individual oocytes were then lysed by incubating in 0.5 ml 10% SDS (v/v) for 1 h at 22°C. Radioactivity was determined via scintillation counting after addition of 4 mL of scintillation cocktail.

### Generation of *ERDL4* overexpressing plants

The full-length coding region of *ERDL4* was amplified with primers *ERDL4_Fwd* (5’-ggg gac aag ttt gta caa aaa agc agg ctt aAT GAG TTT TAG GGA TGA TAA TAC G −3’) and *ERDL4_Rev* (5’-ggg gac cac ttt gta caa gaa agc tgg gta TCA TCT GAA CAA AGC TTG GAT CTC −3’). The purified PCR product was cloned via BP clonase (ThermoFisher, Carlsbad, CA, USA) into pDON/Zeo and cloned via LR Clonase (ThermoFisher, Carlsbad, CA, USA) into the pK2GW7 Gateway vector (Karimi et al., 2002) carrying a *Ca35S* promotor. For generation of *ERDL4-*overexpression lines in tst1/2 double knockout background, ERDL4 coding sequence was amplified with primers harboring *Xho* 1 and *BamH1* restriction enzyme sites. *ERDL4_Xho_s:* 5’ - ctc gag aac aAT GAG TTT TAG GGA TGA TAA TAC G and *ERDL4_Bam_as*: 5’ - gga tcc TCA TCT GAA CAA AGC TTG GAT CTC. Amplified fragment was inserted in pBSK vector (digested with *Eco*RV), cleaved again with *Xho1* and *Bam*H1 and inserted in pHannibal vector (digested with corresponding enzymes). The construct was subsequently digested with *Not1* and cloned into pART27 expression vector (digested with *Not1). Agrobacterium tumefaciens* strain GV3101 was used for transformation. Arabidopsis plants were transformed via floral dip method (Clough and Bent, 1998). Positive lines were screened by Kanamycin antibiotic resistance and RT-PCR was used further for selection of strongest lines.

### Identification of T-DNA knockout lines

T-DNA insertion lines SALK_116041 (insertion at −34 bp, 5’ UTR) and GABI_600B02 (insertion at +2281 bp, exon 14) were provided by the Nottingham Arabidopsis Stock Centre (http://nasc.nott.ac.uk/). T-DNA insertion was confirmed with T-DNA specific primers using genomic DNA. ERDL4 transcript level was checked in both lines with cDNA via PCR and RT-PCR.

### RNA isolation, cDNA synthesis and quantitative Real-Time PCR

RNA isolation was carried out using the NucleoSpin® RNA Plant Kit (Macherey-Nagel) according to the manufacturer’s instructions. 50 mg of fresh plant material was used for this purpose. The RNA was eluted from the column with 40 μl of RNase-free ddH_2_O and the concentration was determined with NanoPhotometer® N50 (Implen). Preparation of cDNA was accomplished by means of qScriptTM cDNA Synthesis Kit (Quantabio) according to manufacturer’s instructions. 1 μg RNA was used for each sample. The resulting cDNA was then used, after dilution to a factor of (1:10), for expression analysis via qRT-PCR. Primers used for the expression analysis are given in Supplemental table S1. Transcripts were normalized using *AtAct2* (*At3G18780*) as reference gene.

### Generation of RNASeq data and analysis

WT and overexpression mutants were grown on soil for 6 weeks. Samples for RNA sequencing were harvested at midday and stored in liquid nitrogen prior to freezing at −80°C. For RNA sequencing of the sugar induced samples, leaf discs from WT plants were incubated in 100mM sugars (Mannitol, Glucose and Fructose) overnight. Leaf discs were harvested and stored at −80°C. Preparation of RNA was done as described above. RNA sequencing was done with Novogene (UK). For correlation analysis Microsoft Excel was used. Heat maps were generated using Orange data mining software (https://orangedatamining.com) with its bioinformatics suite. Volcano plots were generated via online app VolcaNoseR (Goedhart & Luijsterburg, 2020). Custom venn diagrams were prepared using an online web tool (https://bioinformatics.psb.ugent.be/web_tools/Venn/).

### Metabolite quantification and phloem exudate analysis

Extraction of soluble sugars was done with 1ml of 80% Ethanol added to 100 mg of frozen finely ground plant material at 80°C for 30 min. The extracts were collected and evaporated in a vacufuge concentrator (Eppendorf, Hamburg, Germany). Resulting pellets were dissolved in ddH_2_O and quantified using NADP-coupled enzymatic test (Stitt et al., 1989) via 96-well plate reader (Tecan Infinite 200; Tecan Group).

Phloem exudates were collected according to the protocol from (Xu et al., 2019) with few modifications. Source leaves of the same developmental stage from 6 weeks old well-watered plants, were cut with a sharp razor blade at the base of the petiole. Petioles were dipped in 20mM K2-EDTA solution. Leaves of the same line were stacked and the petioles were recut while dipped in the EDTA solution. Leaves were then transferred to 1.5 ml Eppendorf tubes with 500 ul of EDTA solution. Exudates were collected for 6 hours in a dark and humid chamber. Leaves were weighed and exudates were dried in a vacuum concentrator. For sugar quantification pellets were solubilized in 100 ul of ddH_2_O.

### Seed and silique analysis

For seed analysis plants were grown in standard conditions. Completely mature and air-dried seeds were harvested at the same time point for all the lines. For seed weight, 1000 seeds were counted and weighed gravimetrically. Lipid quantification was done according to previously optimized protocols (Reiser et al., 2004). 100 mg of seeds were homogenized in a mortar with liquid nitrogen. 1.5 ml of isopropanol was added to the homogenate and seeds were further homogenized. It was then transferred to 1.5 ml Eppendorf tubes and incubated at 4°C for 12 hours at 100 rpm. Samples were centrifuged at 13,000 g for 10 minutes. Subsequently, supernatant was transferred to weighed 1.5 ml Eppendorf tubes and allowed to evaporate at 60°C for 8 hours. The remaining lipid pellet was quantified gravimetrically. Inflorescence lengths were recorded from day 1 of bolting until they reached their maximum height.

### Electrical conductivity assays

Electrolyte leakage analysis was conducted according to the protocol described by (Sukumaran and Weiser, 1972; Ristic and Ashworth, 1993). A leaf of the same developmental stage, from 3 weeks of cold acclimated plants was dipped in test tubes having 2 ml of deionized water and placed in a cryostat. Ice crystallization was initiated at −2°C by touching the tubes with a wire frozen in liquid nitrogen. Following equilibration for 1 hour, the temperature was reduced at a rate of 2°C/hour and samples were collected at each decrement. Collected samples were thawed on ice overnight. 2 ml of deionized water was added to the samples and ions were extracted by shaking the samples gently for 2 hours. The initial conductivity was measured using a conductivity meter (WTW LF521; Wissenschaftlich-Technische Werkstätten). The samples were then boiled for 2 hours, and ions were re-extracted by shaking for 2 hours. Final conductivity was then measured and LT50% was calculated statistically.

### Non-aqueous fractionation

Plant material for subcellular sugar analysis was grown under short day conditions (10 h light / 14 h dark regime). Leaf material was harvested at middays, snap-frozen and ground to a fine powder under constant supply of liquid nitrogen. The frozen and ground tissue powder was freeze-dried, and 15 to 20 mg were used for subsequent NAF analysis as described earlier (Fürtauer *et al*., 2016). In brief, mixtures of tetrachlorethylen and n-heptane were prepared with total densities ranging between 1.35 and 1.55 g cm^-3^. Freeze dried leaf tissue powder was suspended in the solvent of lowest density, sonicated, filtered and fractionated consecutively by centrifugation. Supernatants were transferred to new tubes and used for maker enzyme activity measurements as well as for determination of sugar amounts. Finally, marker enzyme activities of vacuole, cytosol and chloroplast were correlated to sugar abundance following the protocol described before (Fürtauer *et al*., 2016).

## Acknowledgements

The authors would like to thank Ralf Pennther-Hager (University of Kaiserslautern) for plant growth and Verena Müller and Ruth Wartenberg (University of Kaiserslautern) for excellent technical assistance. We would like to thank Patrick A.W. Klemens for supporting of this project in its initial phase and generation of ERDL4-GFP lines. This research was funded by a DAAD grant to AK (Graduate School Scholarship Program 2015 (57145465). HEN and JSS received additional funding from DFG program RTG2737 (“Stressistance”). HEN received additional funding from the priority program “BioComp 3.0” of the Research initiative of the state Rhineland-Palatine. Research in the lab of HEN and TN was funded by DFG, TR175, Projects B03 and D03, respectively.

## Author contributions

AK generated or characterized plant lines and performed most of the experiments. JC, CP and JS-S conducted oocyte uptake assays. IK conducted cold acclimation kinetics. TN, LF and AnK conducted NAF analysis. BP conceived the study and analyzed RNASeq data. HEN and BP wrote the paper.

## References

Aguero T, Newman K, King ML (2018) Microinjection of Xenopus Oocytes. Cold Spring Harb Protoc 2018

Alonso JM, Stepanova AN, Leisse TJ, Kim CJ, Chen H, Shinn P, Stevenson DK, Zimmerman J, Barajas P, Cheuk R, et al. (2003) Genome-wide insertional mutagenesis of Arabidopsis thaliana. Science 301: 653–657

Aluri, S., Büttner, M. (2007). Identification and functional expression of the Arabidopsis thaliana vacuolar glucose transporter 1 and its role in seed germination and flowering. Proc Natl Acad Sci USA 104: 2537–2542

Aubry E, Hoffmann B, Vilaine F, Gilard F, Klemens PAW, Guérard F, Gakière B, Neuhaus HE, Bellini C, Dinant S, Le Hir R (2022) A vacuolar hexose transport is required for xylem development in the inflorescence stem. Plant Physiol. 188: 1229–1247

Chardon F, Bedu M, Calenge F, Klemens PA, Spinner L, Clement G, Chietera G, Leran S, Ferrand M, Lacombe B, Loudet O, Dinant S, Bellini C, Neuhaus HE, Daniel-Vedele F, Krapp A (2013) Leaf fructose content is controlled by the vacuolar transporter SWEET17 in Arabidopsis. Curr Biol 23: 697–702

Chen HY, Huh JH, Yu YC, Ho LH, Chen LQ, Tholl D, Frommer WB, Guo WJ (2015) The Arabidopsis vacuolar sugar transporter SWEET2 limits carbon sequestration from roots and restricts Pythium infection. Plant J 83: 1046–1058

Chen Q, Xu X, Xu D, Zhang H, Zhang C, Li G (2019) WRK18 and WRKY53 coordinate with HISTONE ACETYLTRANSFERASE1 to regulate rapid responses to sugar. Plant Physiol. 180: 2212–2226.

Cheng J, Wen S, Xiao S, Lu B, Ma M, Bie Z (2017) Overexpression of the tonoplast sugar transporter CmTST2 in melon fruit increases sugar accumulation. J Exp Bot 69: 511–523

Cho Y-H, Yoo S-D (2011) Signaling role of fructose mediated by FINS1/FBP in Arabidopsis thaliana. PLoS Genetics 7: e1001263–e1001263

Cho JI, Burla B, Lee DW, Ryoo N, Hong SK, Kim HB, Eom JS, Choi SB, Cho MH, Bhoo SH, Hahn TR, Neuhaus HE, Martinoia E, Jeon JS (2010) Expression analysis and functional characterization of the monosaccharide transporters, OsTMTs, involving vacuolar sugar transport in rice (Oryza sativa). New Phytol. 186: 657–668

Deng J, Yang X, Sun W, Miao Y, He L, Zhang X (2020) The calcium sensor CBL2 and its interacting kinase CIPK6 are involved in plant sugar homeostasis via interacting with tonoplast sugar transporter TST2. Plant Physiol. 183: 236–249.

Dhungana SR, Braun DM (2021) Sugar transporters in grasses: Function and modulation in source and storage tissues. J. Plant Physiol. 266: 153541

Eom JS, Chen LQ, Sosso D, Julius BT, Lin IW, Qu XQ, Braun MD, Frommer WB (2015) SWEETs, transporters for intracellular and intercellular sugar translocation. Curr Op Plant Biol. 25: 53–62

Furuichi T, Mori IC, Takahashi K, Muto S (2001) Sugar-induced increase in cytosolic Ca2+ in Arabidopsis thaliana whole plants. Plant Cell Physiol 42: 1149–1155

Fürtauer L, Weckwerth W, Nägele T (2016) A benchtop fractionation procedure for subcellular analysis of the plant metabolome. Front Plant Sc 7: 1912

Fürtauer L, Küstner L, Weckwerth W, Heyer AG, Nägele T (2019) Resolving subcellular plant metabolism. The Plant Journal 100: 438–455

Guo WJ, Nagy R, Chen HY, Pfrunder S, Yu YC, Santelia D, Frommer WB, Martinoia E (2014) SWEET17, a facilitative transporter, mediates fructose transport across the tonoplast of Arabidopsis roots and leaves. Plant Physiol 164: 777–789

Hedrich R, Sauer N, Neuhaus HE (2015) Sugar transport across the plant vacuolar membrane: nature and regulation of carrier proteins. Curr. Opin. Plant Biol. 25: 63–70

Henderson PJF (1991) Sugar transport proteins. Curr. Opin. Struc. Biol. 1: 590–601

Ho LH, Rode R, Siegel M, Reinhardt F, Neuhaus HE, Yvin JC, Pluchon S, Hosseini SA, Pommerrenig B (2020) Potassium application boosts photosynthesis and sorbitol biosynthesis and accelerates cold acclimation of common plantain (Plantago major L.) Plants 9: 1259

Hu D-G, Sun C-H, Zhang Q-Y, An J-P, You C-X, Hao Y-J (2016) Glucose sensor MdHXK1 phosphorylates and stabilizes MdbHLH3 to promote anthocyanin biosynthesis in apple. PLoS Gen. 12:e1006273

Hunter K (2020) CBL2-CIPK6-TST2-mediated regulation of sugar homeostasis. Plant Physiol. 183: 21–22

Jang JC, Sheen J (1994) Sugar sensing in higher plants. Plant Cell 6: 1665–1679

Jänkänpää HJ, Mishra Y, Schröder WP, Jansson S (2012) Metabolic profiling reveals metabolic shifts in Arabidopsis plants grown under different light conditions. Plant, Cell & Environ. 35: 1824–1836

Julius BT, Leach KA, Tran TM, Mertz RA, Braun DM (2017) Sugar transporters in plants: new insights and discoveries. Plant Cell Physiol. 58: 1442–1460

Jung B, Ludewig F, Schulz A, Meißner G, Wöstefeld N, Flügge UI, Pommerrenig B, Wirsching P, Sauer N, Koch W, Sommer F, Mühlhaus T, Schroda M, Cuin TA, Graus D, Marten I, Hedrich R, Neuhaus HE (2015) Identification of the transporter responsible for sucrose accumulation in sugar beet taproots. Nature Plants 1: Article number: 14001

Keller I, Rodrigues CM, Neuhaus HE, Pommerrenig B (2021) Improved resource allocation and stabilization of yield under abiotic stress. J. Plant Physiol. 257: 153336

Kiyosue T, Abe H, Yamaguchi-Shinozaki K, Shinozaki K (1998) ERD6, a cDNA clone for an early dehydration-induced gene of Arabidopsis, encodes a putative sugar transporter. Biochim Biophys Acta 1370: 187–191

Klemens PA, Patzke K, Deitmer J, Spinner L, Le HR, Bellini C, Bedu M, Chardon F, Krapp A, Neuhaus HE (2013) Overexpression of the vacuolar sugar carrier AtSWEET16 modifies germination, growth, and stress tolerance in Arabidopsis. Plant Physiol 163: 1338–1352

Klemens PA, Patzke K, Trentmann O, Poschet G, Büttner M, Schulz A, Marten I, Hedrich R, Neuhaus HE (2014) Overexpression of a proton-coupled vacuolar glucose exporter impairs freezing tolerance and seed germination. New Phytol. 202: 188–197

Komarova NY, Meier S, Meier A, Suter-Grotemeyer M, Rentsch D (2012) Determinants for Arabidopsis peptide transporter targeting to the tonoplast or plasma membrane. Traffic 13: 1090–1105

Krueger S, Giavalisco P, Krall L, Steinhauser MC, Bussis D, Usadel B, Flugge UI, Fernie AR, Willmitzer L, Steinhauser D (2011) A topological map of the compartmentalized Arabidopsis thaliana leaf metabolome. PLoS.One. 6: e17806

Lecompte F, Nicot PC, Ripoll J, Abro MA, Raimbault AK, Lopez-Lauri F, Bertin N (2017) Reduced susceptibility of tomato stem to the necrotrophic fungus Botrytis cinerea is associated with a specific adjustment of fructose content in the host sugar pool. Ann. Bot 119: 931–943

Le Hir R, Spinner L, Klemens PAW, Chakraborti D, de Marco F, Vilaine F, Wolff N, Lemoine R, Porcheron B, Géry C, Téoulé E, Chabout S, Mouille G, Neuhaus HE, Dinant S, Bellini C (2015) Disruption of the sugar transporters AtSWEET11 and AtSWEET12 affects vascular development and freezing tolerance in Arabidopsis. Mol. Plant 8: 1687–1690

Lemoine R, La CS, Atanassova R, Dedaldechamp F, Allario T, Pourtau N, Bonnemain JL, Laloi M, Coutos-Thevenot P, Maurousset L, Faucher M, Girousse C, Lemonnier P, Parrilla J, Durand M (2013) Source-to-sink transport of sugar and regulation by environmental factors. Front Plant Sci 4: 272

Li P, Wind JJ, Shi X, Zhang H, Hanson J, Smeekens SC, Teng S (2011) Fructose sensitivity is suppressed in Arabidopsis by the transcription factor ANAC089 lacking the membrane-bound domain. Proc. Natl. Acad. Sci. 108: 3436–3441

Luo QJ, Mittal A, Jia F, Rock CD (2012). An autoregulatory feedback loop involving PAP1 and TAS4 in response to sugars in Arabidopsis. Plant Mol Biol. 80: 117–129

Martinoia E, Meyer S, De Angeli A, Nagy R (2012) Vacuolar transporters in their physiological context. Ann. Rev. Plant Biol. 63: 163–213

McKown R, Kuroki G, Warren G (1996). Cold responses of Arabidopsis mutants impaired in freezing tolerance. J. Exp. Bot. 47: 1919–1925

Nägele T, Heyer AG (2013) Approximating subcellular organisation of carbohydrate metabolism during cold acclimation in different natural accessions of Arabidopsis thaliana. New Phytol. 198: 777–787

Nägele T, Henkel S, Hörmiller I, Sauter T, Sawodny O, Ederer M, Heyer AG (2010) Mathematical Modeling of the Central Carbohydrate Metabolism in Arabidopsis Reveals a Substantial Regulatory Influence of Vacuolar Invertase on Whole Plant Carbon Metabolism, Plant Physiol. 153: 260–272

Nørholm MHH, Nour-Eldin HH, Brodersen P, Mundy J, Halkier BA (2006) Expression of the Arabidopsis high-affinity hexose transporter STP13 correlates with programmed cell death. FEBSL. 580: 2381–2387

Okooboh GO, Haferkamp I, Valifard M, Pommerrenig B, Kelly A, Feussner I, Neuhaus HE (2022) Overexpression of the vacuolar sugar importer BvTST1 from sugar beet in Camelina improves seed properties and leads to altered root characteristics. Physiol. Plant. 174: e13653.

Patzke K, Prananingrum P, Klemens PAW, Trentmann O, Martins Rodrigues C, Keller I, Fernie AR, Geigenberger P, Bölter B, Lehmann M, Schmitz-Esser S, Pommerrenig B, Haferkamp I, Neuhaus HE (2019) The plastidic sugar transporter pSuT influences flowering and affects cold responses. Plant Physiol. 179: 569–587

Pertl-Obermeyer H, Trentmann O, Duscha K, Neuhaus HE, Schulze WX (2016) Quantitation of vacuolar sugar transporter abundance changes using QconCAT synthtetic peptides. Front. Plant Sc. 7: 411.

Pommerrenig B, Ludewig F, Cvetkovic J, Trentmann O, Klemens PAW, Neuhaus HE (2018) In concert: orchestrated changes in carbohydrate homeostasis are critical for plant abiotic stress tolerance. Plant Cell Physiol. 59: 1290–1299

Pommerrenig B, Eggert K, Bienert GP (2019) Boron deficiency effects on sugar, ionome, and phytohormone profiles of vascular and non-vascular leaf tissues of common plantain (Plantago major L.) Int. J. Mol. Sci. 20: 3882

Pommerrenig B, Müdsam C, Kischka D, Neuhaus HE (2020) Treat and trick: common regulation and manipulation of sugar transporters during sink establishment by the plant and the pathogen. J. Exp. Bot. 71: 3930–3940

Poschet G, Hannich B, Raab S, Jungkunz I, Klemens PA, Krüger S, Wic S, Neuhaus HE, Büttner M (2011) A novel Arabidopsis vacuolar glucose exporter is involved in cellular sugar homeostasis and affects the composition of seed storage compounds. Plant Physiol 157: 16641676

Remy E, Cabrito TR, Batista RA, Hussein MA, Teixeira MC, Athanasiadis A, Sá-Correia I, Duque P (2014) Intron retention in the 5’ UTR of the novel ZIF2 transporter enhances translation to promote zinc tolerance in Arabidopsis. PLoS genetics 10: e1004375.

Rodrigues CM, Müdsam C, Keller I, Zierer W, Czarnecki O, Corral JM, Reinhardt F, Nieberl P, Fiedler-Wiechers K, Sommer F, Schroda M, Mühlhaus T, Harms K, Flügge UI, Sonnewald U, Koch W, Ludewig F, Neuhaus HE, Pommerrenig B (2020). Vernalization alters sink and source identities and reverses phloem translocation from taproots to shoots in sugar beet. Plant Cell 32: 3206–3223

Rolland F, Baena-Gonzalez E, Sheen J (2006) Sugar sensing and signaling in plants: conserved and novel mechanisms. Annu. Rev. Plant Biol. 57: 675–709

Rottmann T, Klebl F, Schneider S, Kischka D, Rüscher D, Sauer N, Stadler R (2018) Sugar transporter STP7 specificity for l-arabinose and d-xylose contrasts with the typical hexose transporters STP8 and STP12. Plant Physiol. 176: 2330–2350

Ruan YL (2014) Sucrose metabolism: gateway to diverse carbon use and sugar signaling. Ann. Rev. Plant Biol. 65: 33–67

Slawinksi L, Israel A, Paillot C, Thibault F, Cordaux R, Atanassova R, Dédaldéchamp F, Laloi M (2021a) Early response to dehydration six-like transporter family: early origin in streptophytes and evolution in land plants. Front Plant Sci. 12: 681929

Slawinski L, Israel A, Artault C, Thibault F, Atanassova R, Laloi M, Dédaldéchamp F (2021b) Responsiveness of early response to dehydration six-like transporter genes to water deficit in Arabidopsis thaliana leaves. Front. Plant Sci. 12: 708876

Schofield RA, Bi YM, Kant S, Rothstein SJ (2009) Over-expression of STP13, a hexose transporter, improves plant growth and nitrogen use in Arabidopsis thaliana seedlings. Plant Cell Environ. 32: 271–285

Schneider S, Hulpke S, Schulz A, Yaron I, Holl J, Imlau A, Schmitt B, Batz S, Wolf S, Hedrich R, Sauer N (2012) Vacuoles release sucrose via tonoplast-localised SUC4-type transporters. Plant Biol. 14: 325–336

Schulz A, Beyhl D, Marten I, Wormit A, Neuhaus E, Poschet G, Büttner M, Schneider S, Sauer N, Hedrich R (2011) Proton-driven sucrose symport and antiport are provided by the vacuolar transporters SUC4 and TMT1/2. Plant J. 68: 129–136

Schulze WX, Schneider T, Starck S, Martinoia E, Trentmann O (2012) Cold acclimation induces changes in Arabidopsis tonoplast protein abundance and activity and alters phosphorylation of tonoplast monosaccharide transporters. Plant J. 69: 529–541

Smeekens S (1998) Sugar regulation of gene expression in plants. Curr Op plant biol 1: 230–234.

Smith AM, Stitt M (2007) Coordination of carbon supply and plant growth. Plant, Cell Environ. 30: 1126–1149

Smith AM, Zeeman SC (2020) Starch: a flexible, adaptable carbon store coupled to plant growth. Ann. Rev. Plant Biol. 71: 217–245

Stitt M, Hurry V (2002) A plant for all seasons: alterations in photosynthetic carbon metabolism during cold acclimation in Arabidopsis. Curr. Opin. Plant Biol. 5: 199–206

Stitt M, Wirtz W, Gerhardt R, Heldt HW, Spencer CA, Walker D, Foyer CH (1985) A comparative study of metabolite levels in plant leaf material in the dark. Planta 166: 354–364

Strand Å, Hurry V, Gustafsson P, Gardeström P (1997) Development of Arabidopsis thaliana leaves at low temperatures releases the suppression of photosynthesis and photosynthetic gene expression despite the accumulation of soluble carbohydrates. Plant J, 12: 605–614

Sukumaran, N. P., & Weiser, C. J. (1972) An Excised Leaflet Test for Evaluating Potato Frost Tolerance. HortScience 7: 467–468

Sun L, Sui X, Lucas WJ, Li Y, Feng S, Ma S, Fan J, Gao L, Zhang, Z (2019) Down-regulation of the sucrose transporter CsSUT1 causes male sterility by altering carbohydrate supply. Plant Physiol. 180: 986–997

Truernit E, Sauer N (1995) The promoter of the Arabidopsis thaliana SUC2 sucrose-H+ symporter gene directs expression of beta-glucuronidase to the phloem: evidence for phloem loading and unloading by SUC2. Planta 196: 564–570

Valifard M, Le-Hir R, Müller J, Scheuring D, Neuhaus HE, Pommerrenig B (2021) The Vacuolar fructose transporter SWEET17 is critical for root development and drought tolerance. Plant Physiology 187: 2716–2730

Volkert K, Debast S, Voll LM, Voll H, Schießl I, Hofmann J, Schneider S, Börnke F (2014) Loss of the two major leaf isoforms of sucrose-phosphate synthase in Arabidopsis thaliana limits sucrose synthesis and nocturnal starch degradation but does not alter carbon partitioning during photosynthesis. J. Exp. Bot. 65: 5217–5229

Vu DP, Martins Rodrigues C, Jung B, Meissner G, Klemens PAW, Holtgräwe D, Fürtauer L, Nägele T, Nieberl P, Pommerrenig B, Neuhaus HE (2020). Vacuolar sucrose homeostasis is critical for plant development, seed properties, and night-time survival in Arabidopsis. J Ex Bot 71: 4930–4943

Wieczorke R, Krampe S, Weierstall T, Freidel K, Hollenberg CP, Boles E (1999). Concurrent knock-out of at least 20 transporter genes is required to block uptake of hexoses in Saccharomyces cerevisiae. FEBS letters, 464: 123–128

Wingenter K, Schulz A, Wormit A, Wic S, Trentmann O, Hörmiller II, Heyer AG, Marten I, Hedrich R, Neuhaus HE (2010) Increased activity of the vacuolar monosaccharide transporter TMT1 alters cellular sugar partitioning, sugar signalling and seed yield in Arabidopsis. Plant Physiol. 154: 665–677

Wingenter K, Trentmann O, Winschuh I, Hörmiller II, Heyer AG, Reinders J, Schulz A, Geiger D, Hedrich R, Neuhaus HE (2011) A member of the mitogen-activated protein 3-kinase family is involved in the regulation of plant vacuolar glucose uptake. Plant J 68: 890–900

Wolfenstetter S, Wirsching P, Dotzauer D, Schneider S, Sauer N (2012) Routes to the tonoplast: The sorting of tonoplast transporters in Arabidopsis mesophyll protoplasts. Plant Cell 24: 215–232.

Wormit A, Trentmann O, Feifer I, Lohr C, Tjaden J, Meyer S, Schmidt U, Martinoia E, Neuhaus HE (2006) Molecular identification and physiological characterization of a novel monosaccharide transporter from Arabidopsis involved in vacuolar sugar transport. Plant Cell 18: 3476–3490

Yamada K, Osakabe Y, Mizoi J, Nakashima K, Fujita Y, Shinozaki K, Yamaguchi-Shinozaki K (2010) Functional analysis of an Arabidopsis thaliana abiotic stress-inducible facilitated diffusion transporter for monosaccharides. J Biol Chem 285: 1138–1146

Zhong Y, Xie J, Wen S, Wu W, Tan L, Lei M, Shi H, Zhu JK (2020) TPST is involved in fructose regulation of primary root growth in Arabidopsis thaliana. Plant Mol. Biol. 103: 511–525

Zhu L, Li B, Wu L, Li H, Wang Z, Wei X, Ma B, Zhang Y, Ma F, Ruan Y-L, Li M (2021) MdERDL6-mediated glucose efflux to the cytosol promotes sugar accumulation in the vacuole through up-regulating TSTs in apple and tomato. Proc. Natl. Acad. Sci. 118: e2022788118

