## Supplemental Information for "The vacuolar sugar transporter *Early Response to Dehydration 6-Like4* regulates fructose signaling and plant growth"

A

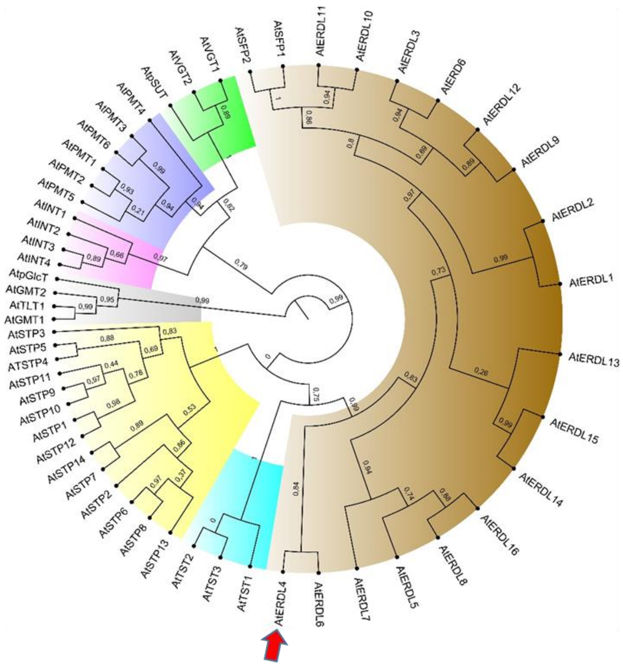


B

001 MSFRDDNTEE GRNDLRRPFL HTGSWYRMGS RQSSMLESSQ VIRDSSISVL

051 ACVLIVALGP IQFGFTCGYS SPTQAAITKD LGLTVSEYSV FGSLSNVGAM

101 VGAIASGQIA EYVGRKGSLM IAAIPNIIGW LSISFAKDTS FLYMGRLLEG

151 FGVGIISYTV PVYIAEIAPQ TMRGALGSVN QLSVTIGIML AYLLGLFVPW

201 RILAVLGVLP CTLLIPGLFF IPESPRWLAK MGLTDDFETS LQVLRGFETD

251 ITVEVNEIKR SVASSSKRSA VRFVDLKRRR YYFPLMVGIG LLALQQLGGI

301 NGVLFYSSTI FESAGVTSSN VATFGVGVVQ VVATGIATWL VDKAGRRLLL

351 M**I**SSIGMTIS LVIVAVAFYL KEFVSPDSNM YNILSMVSVV GVVAMVISCS

401 LGMGPIPWLI MSEILPVNIK GLAGSIATLL NWFVSWLVTM TANMLLAWSS

451 GGTFTLYALV CGFTVVFVSL WVPETKGKTL EEIQALFR

**Figure S1. Phylogeny of Arabidopsis MST proteins and amino acid sequence of ERDL4**

**A)** Phylogeny of Arabidopsis MST proteins with focus on the ERDL subgroup (brown shading). The red arrowhead points towards ERDL4. The members of the ERDL group have the following identifiers: ERD6: At1g08930; ERDL1:At1g08890; ERDL2: At1g08900; ERDL3: At1g08920; ERDL4: At1g19450; ERDL5: At1g54730; ERDL6: At1g75220; ERDL7: At2g48020; ERDL8:At3g05150; ERDL9: At3g05155; ERDL10: At3g05160; ERDL11: At3g05165; ERDL12: At3g05400; ERDL13:At3g20460; ERDL14:At4g04750; ERDL15:At4g04760; ERDL16: At5g18840; SFP1: At5g27350; SFP2: At5g27360 **B)** Amino acid sequence of the ERDL4 protein. Predicted transmembrane domains (TmPred) are highlighted in grey. The bold isoleucine residue highlighted in yellow at position 352 marks the putative end of the native protein sequence in *erdl4-2* mutants due to the presence of the T-DNA insertion. Underlined stretches indicate the conserved sugar transport motif (Sugar_tr; PF00083).

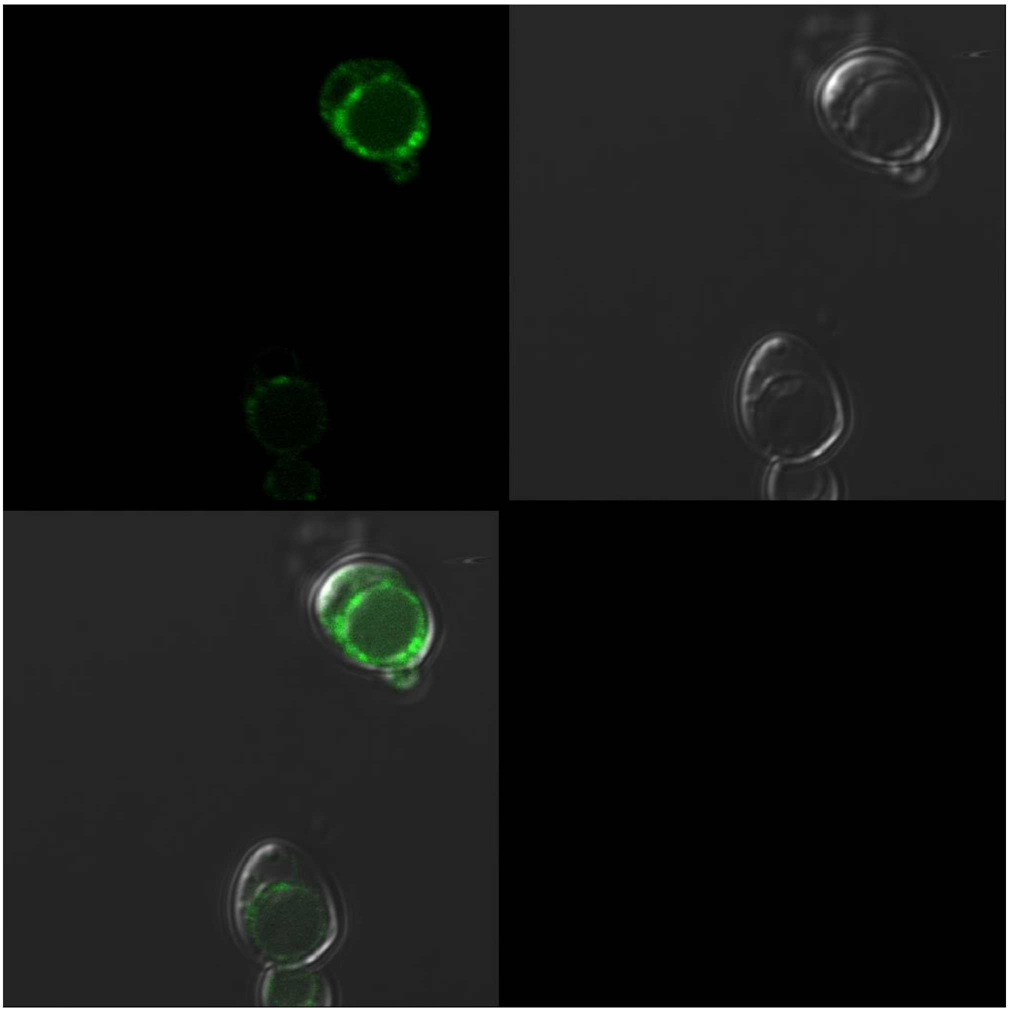

**Supplementary Figure S2.** Localization of ERDL4-GFP in yeast. Green GFP-derived fluorescence was detected in ring-like structures that could be assigned to yeast vacuoles and other internal membranes.

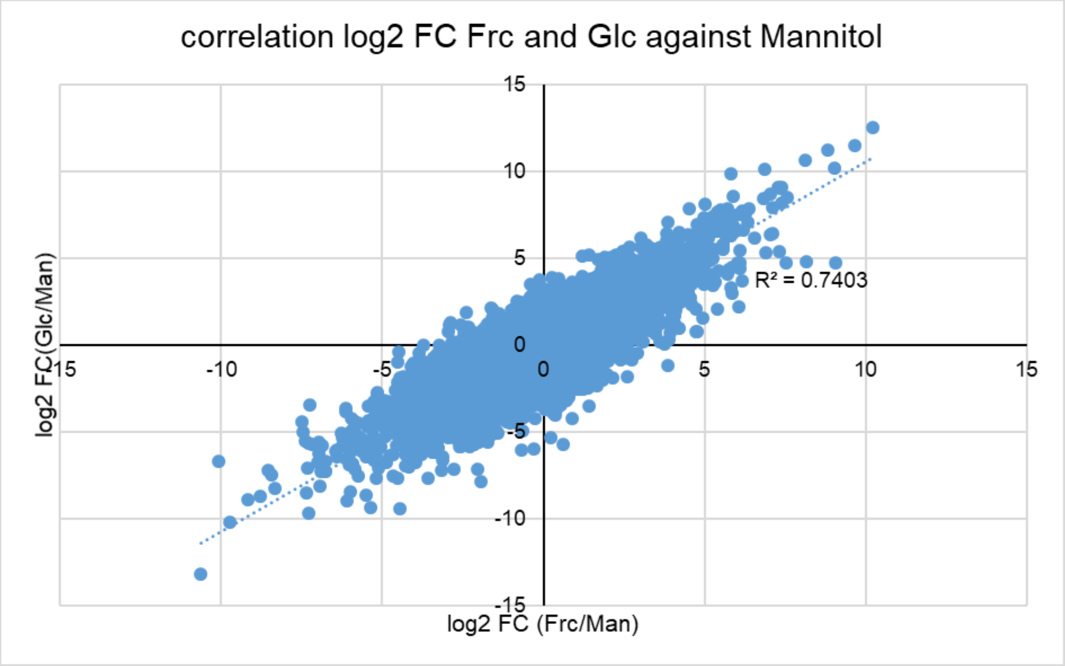


**Supplementary Figure S3**. Correlation between the log2FCs of glucose-regulated genes (on y-axis) and of fructose-regulated genes (on x-axis).

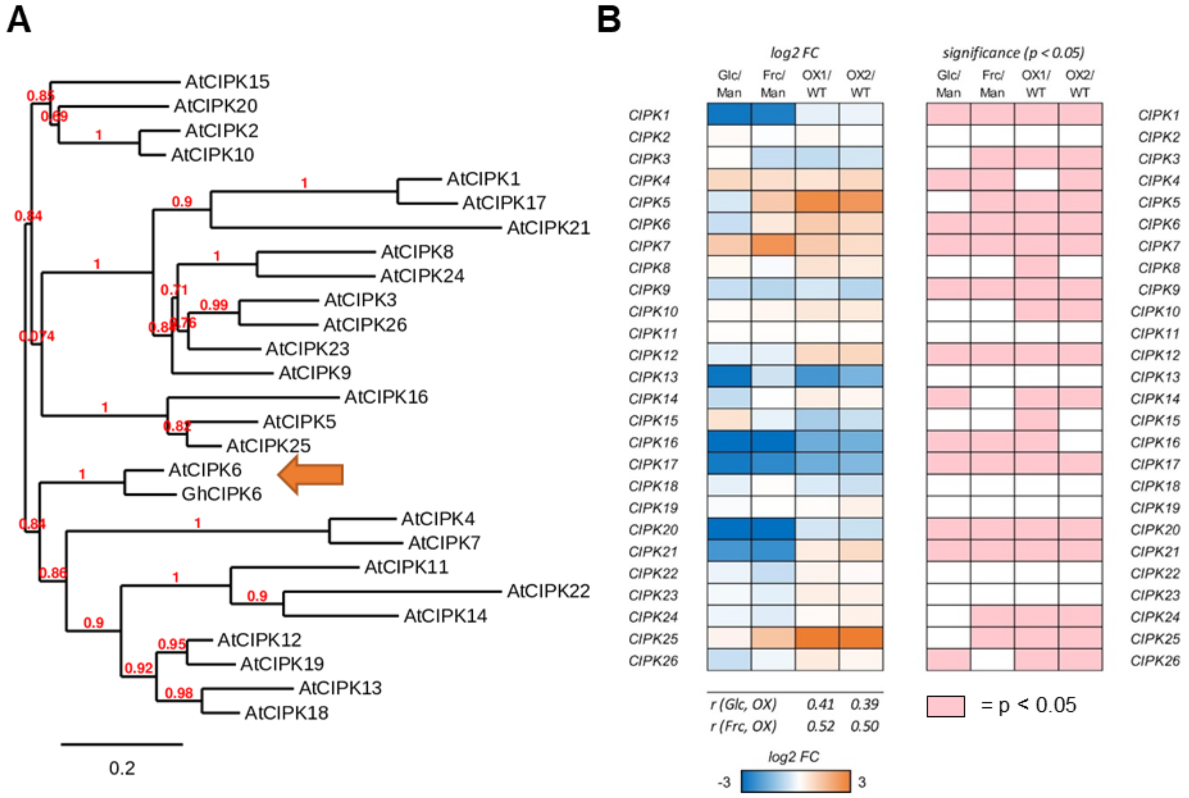


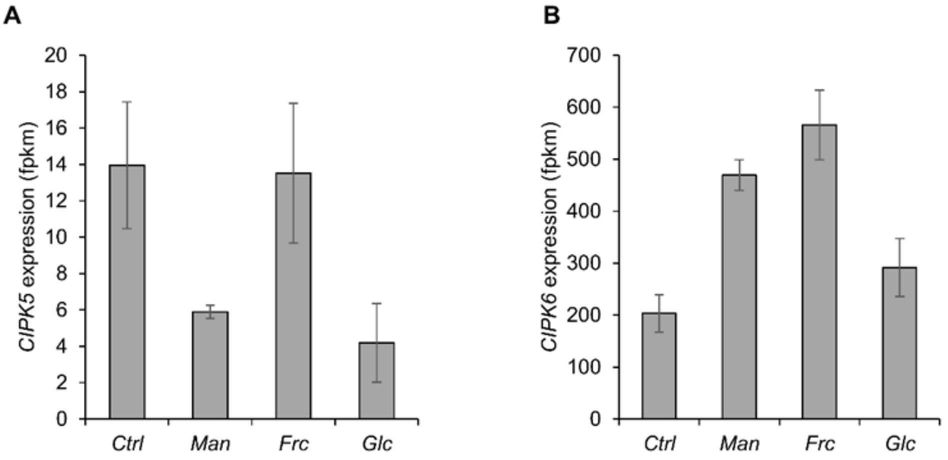


**Supplementary Figure S4. A)** Phylogenetic sequence comparison of Arabidopsis CIPK proteins with GhCIPK6 B) Heat map representation of *At*CIPK expression (based on fold change of fpkm vlues) of glucose (glc) or fructose (frc) incubated leaf discs (relative to mannitol as osmotic control) in *35S-ERDL4 #1* (OX1) and *35S-ERDL4 #2* (OX2) plants (relative to WT). C, D) Absolute expression of CIPK5 and CIPK6

### Table S1. Primers used in this study

### Oligonucleotides used for RT-qPCR

| gene | gene ID | Forward (5´→ 3´) | Reverse (5´→ 3´) |
|---|---|---|---|
| ***AtActin2*** | *At3G18780* | ACGGTAACATTGTGCTCAGTGGTG | CTTGGAGATCCACATCTGCTGGA |
| ***AtERDL4*** | *At1G19450* | TGGAGTTGGCGTTGTTCAGGTAG | GGATAAAGCAGGTCGTCGGCTT |
| ***AtERDL6*** | *At1G75220* | GGTCGTCGGCTTCTGCTTACTATC | GCCTTGTAATTGTTGCAGCTGCT |
| ***AtTST1*** | *At1G20840* | TTGCCGGCGAATTCTACTAAAGAG | CAAGGATGGGACAATGCCACCA |
| ***AtTST2*** | *At4G35300* | CATGGATCTTTCTGGTCGAAGGAC | GATAAGACCGCGTGCACAATGC |
| ***AtVGT1*** | *At3G03090* | TCGAGGCAAATGCTTGAAAGCTC | CCAACAGATAACTGGGCAACCAA |
| ***AtVGT2*** | *At5G17010* | ATGCACCATCAATACTGCAGACC | CAGGTGATGCAACAAGGGTCTCA |
| ***AtVGT3*** | *At5G59250* | ATGCGGGTTCGATTCTTCAGAC | ACAGAGACTCGAGTTGCATCAGC |
| ***AtSweet16*** | *At3G16690* | GAGATGCAAACTCGCGTTCTAGT | GCACACTTCTCGTCGTCACA |
| ***AtSweet17*** | *At4G15920* | AGTGACAACAAAGAGCGTGAAATAC | ACTTAAACCGTTGCTTAAACCAACC |
| ***AtCAB1*** | *At1G29930* | TTACTTGCGCCACACTCTCACC | TTTCCGGTCAAAGCAGGAGAGG |
| ***AtNR1*** | *At1G77760* | CTGAGCTGGCAAATTCCGAAGC | TGCGTGACCAGGTGTTGTAATC |
| ***AtABI4*** | *At2G40220* | GGTCCGTACGGTATCCCTTT | GGTGTTGGAATTGTCCCATC |
| ***AtVIK*** | *At1G14000* | ATGGCTCCTGAAGTATTCAAGC | TCTTGAGAATGTCCAGAAACGACG |

### Primers for genotyping

| primer name | sequence (5´→ 3´) |
|---|---|
| ***Salk_LP*** | ATATGTGAGCGAACATGGGAC |
| ***SALK_LBb1.3*** | ATTTTGCCGATTTCGGAAC |
| ***SALK_RP*** | TTGCAAATGATATCGAAAGCC |
| ***GK_Fwd*** | ATGAGTTTTAGGGATGATAATACGG |
| ***GK_LBP*** | CCCATTTGGACGTGAATGTAGACAC |
| ***GK_Rev*** | TCATCTGAACAAACCTTGGATC |
| ***ERDL4_F*** | ATGAGTTTTAGGGATGATAATAC |
| ***ERDL4_R*** | TCATCTGAACAAAGCTTGGATCTC |
| ***Ef1-Alpha_F*** | GAGACCACCAAGTACTACTGCAC |
| ***EF1-Alpha_R*** | GTTGGTCCCTTGTACCAGTCAAG |

**Supplemental Information 1:** CLUSTAL omega (1.2.4) multiple sequence alignment of AtERDL proteins. Yellow highlights mark putative dileucine-based motifs. Turquoise highlights mark regions belonging to the sugar_tr motif (PF00083).

```
AtERDL14      -----------------------------MA------EESLLPSHTEDVSASPNKSSSLL   25
AtERDL15      -----------------------------MA------EEGLLLP-----ASSTSSSSSLL   20
AtERDL4       ----------------------MSFRDDNTEEGRNDLRRPFLHTGS-WYRMG-SRQSSML   36
AtERDL6       ----------------------MSFRDDN-EEARNDLRRPFIHTGS-WYRMG-SRQSSMM   35
AtERDL5       ------------------------MRGEI-DEANLAPETSLINK----------------   19
AtERDL7       -----------------------------MSKASDAVREPLVD-K---------------   15
AtERDL8       ----------------------METRKDDMEKRNDK-SEPLLLPEN-G------------   24
AtERDL16      ---------------MAIREIKDVERGEIVNKVEDL-GKPFLTHED-D------------   31
AtERDL13      MGDEPLLQKVKIQED--IESVPLLQKVK-IQE---DIESVKGIRVN--------------   40
AtERDL1       -------------------------------MESGSMKTP-LVNNQ--------------   14
AtERDL2       -------------------------------MESERLESH-LLNKQ--------------   14
AtERDL9       -----------------------------MEGENSSIEKGLLLIRK--------------   17
AtERDL12      -----------------------------MEGEN-NMEKGLLLAKK--------------   16
AtERDL10      ------------------------------------MEEGLLRHEN--------------   10
AtERDL11      --------------------------MV-VEEENRSMEEGLLQHQN--------------   19
AtSFP1        ---------------------------M-VMEEGRSIEEGLLQLKN--------------   18
AtSFP2        -----------------MHK------MM-VVEKERSIEERLLQLKN--------------   22
AtERD6        MERQKSMEKGLLRKSLSIRERKFPNEDA-FLESGLSRKSPREVKKP--------------   45
AtERDL3       --------------------MTMSENSR-NL------EAGLLLRKN--------------   19

AtERDL14      SE--ISNASTRPFVLAFTVGSCGALSFGCIVGYTAPTQSSIMKDLNLSIADFSFFGSILT   83
AtERDL15      SE--ISNACTRPFVLAFIVGSCGAFAFGCIIGYSAPTQTSIMKDLNLSIADAIFTIWIDI   78
AtERDL4       ESSQVIRDSSISVLACVLIVALGPIQFGFTCGYSSPTQAAITKDLGLTVSEYSVFGSLSN   96
AtERDL6       GSSQVIRDSSISVLACVLIVALGPIQFGFTCGYSSPTQAAITKDLGLTVSEYSVFGSLSN   95
AtERDL5       ENQDSSATITTTLLLTTFVAVSGSFVFGSAIGYSSPVQSDLTKELNLSVAEYSLFGSILT   79
AtERDL7       NMAGSKPDQPWMVYLSTFVAVCGSFAFGSCAGYSSPAQAAIRNDLSLTIAEFSLFGSLLT   75
AtERDL8       SD--VSEEASWMVYLSTIIAVCGSYEFGTCVGYSAPTQFGIMEELNLSYSQFSVFGSILN   82
AtERDL16      EKESENNESYLMVLFSTFVAVCGSFEFGSCVGYSAPTQSSIRQDLNLSLAEFSMFGSILT   91
AtERDL13      NDGEEDGPVTLILLFTTFTALCGTFSYGTAAGFTSPAQTGIMAGLNLSLAEFSFFGAVLT   100
AtERDL1       EEARSSSSITCGLLLSTSVAVTGSFVYGCAMSYSSPAQSKIMEELGLSVADYSFFTSVMT   74
AtERDL2       EEE--ASSFTSGLLLSTSVVVAGSFCYGCAMSYSSPAQSKIMEELGLSVADYSFFTSVMT   72
AtERDL9       E---ESANTTPFLVFTTFIIVSASFSFGVALGHTAGTMASIMEDLDLSITQFSVFGSLLT   74
AtERDL12      E---DSANTTPLLIFSTFIIVSASFTFGAAIGYTADTMSSIMSDLDLSLAQFSLFGSLST   73
AtERDL10      --DRDDRRITACVILSTFVAVCSSFSYGCANGYTSGAETAIMKELDLSMAQFSAFGSFLN   68
AtERDL11      --DRDDRRITACVILSTFVAVCSAFSYGCAAGYTSGAETAIMKELDLSMAQFSAFGSFLN   77
AtSFP1        KNDDSECRITACVILSTFVAVCGSFSFGVATGYTSGAETGVMKDLDLSIAQFSAFGSFAT   78
AtSFP2        QNDDSECRITACVILSTFIAVCGSFSFGVSLGYTSGAEIGIMKDLDLSIAQFSAFASLST   82
AtERD6        QNDDGECRVTASVFLSTFVAVSGSFCTGCGVGFSSGAQAGITKDLSLSVAEYSMFGSILT   105
AtERDL3       QNDINECRITAVVLFSTFVSVCGSFCFGCAAGYSSVAQTGIINDLGLSVAQYSMFGSIMT   79
                          .         .    *   ..:: .   :   *.*: ::      .

AtERDL14      VGLILGALICGKLADLVGRVYTIWITNILVLIGWLAIAFAKDVRLLDLGRLLQGISVGIS   143
AtERDL15      DG--------GVNPWSINLWETIWITNILFVIGWFAIAFAKGVWLLDLGRLLQGISIGIS   130
AtERDL4       VGAMVGAIASGQIAEYVGRKGSLMIAAIPNIIGWLSISFAKDTSFLYMGRLLEGFGVGII   156
AtERDL6       VGAMVGAIASGQIAEYIGRKGSLMIAAIPNIIGWLCISFAKDTSFLYMGRLLEGFGVGII   155
AtERDL5       IGAMIGAAMSGRIADMIGRRATMGFSEMFCILGWLAIYLSKVAIWLDVGRFLVGYGMGVF   139
AtERDL7       FGAMIGAITSGPIADLVGRKGAMRVSSAFCVVGWLAIIFAKGVVALDLGRLATGYGMGAF   135
AtERDL8       MGAVLGAITSGKISDFIGRKGAMRLSSVISAIGWLIIYLAKGDVPLDFGRFLTGYGCGTL   142
AtERDL16      IGAMLGAVMSGKISDFSGRKGAMRTSACFCITGWLAVFFTKGALLLDVGRFFTGYGIGVF   151
AtERDL13      IGGLVGAAMSGKLADVFGRRGALGVSNSFCMAGWLMIAFSQATWSLDIGRLFLGVAAGVA   160
AtERDL1       LGGMITAAFSGKIAAVIGRRQTMWIADVFCIFGWLAVAFAHDKMLLNIGRGFLGFGVGLI   134
AtERDL2       LGGMITAVFSGKISALVGRRQTMWISDVCCIFGWLAVAFAHDIIMLNTGRLFLGFGVGLI   132
AtERDL9       FGGMIGALFSATIADSFGCKMTLWITEVFCISGWLAIALAKNIIWLDLGRFFVGIGVGLL   134
AtERDL12      FGGMIGAIFSAKAASAFGHKMTLWVADLFCITGWLAISLAKDIIWLDMGRFLVGIGVGLI   133
AtERDL10      LGGAVGALFSGQLAVILGRRRTLWACDLFCIFGWLSIAFAKNVLWLDLGRISLGIGVGLT   128
AtERDL11      VGGAVGALFSGQLAVILGRRRTLWACDFFCVFGWLSIAFAKNVFWLDLGRISLGIGVGLI   137
AtSFP1        LGAAIGALFCGNLAMVIGRRGTMWVSDFLCITGWLSIAFAKEVVLLNFGRIISGIGFGLT   138
AtSFP2        LGAAIGALFSGKMAIILGRRKTMWVSDLLCIIGWFSIAFAKDVMWLNFGRISSGIGLGLI   142
AtERD6        LGGLIGAVFSGKVADVLGRKRTMLFCEFFCITGWLCVALAQNAMWLDCGRLLLGIGVGIF   165
```

```
AtERDL3         FGGMIGAIFSGKVADLMGRKGTMWFAQIFCIFGWVAVALAKDSMWLDIGRLSTGFAVGLL   139
                 *        .      .   ::         **. : :::    *  **   * . *

AtERDL14        SYLGPIYISELAPRNLRGAASSLMQLFVGVGLSAFYALGTAVAWRSLAILGSIPSLVVLP   203
AtERDL15        VYLGPVYITEIAPRNLRGAASSFAQLFAGVGISVFYALGTIVAWRNLAILGCIPSLMVLP   190
AtERDL4         SYTVPVYIAEIAPQTMRGALGSVNQLSVTIGIMLAYLLGLFVPWRILAVLGVLPCTLLIP   216
AtERDL6         SYTVPVYIAEIAPQNMRGGLGSVNQLSVTIGIMLAYLLGLFVPWRILAVLGILPCTLLIP   215
AtERDL5         SFVVPVYIAEITPKGLRGGFTTVHQLLICLGVSVTYLLGSFIGWRILALIGMIPCVVQMM   199
AtERDL7         SYVVPIFIAEIAPKTFRGALTTLNQILICTGVSVSFIIGTLVTWRVLALIGIIPCAASFL   195
AtERDL8         SFVVPVFIAEISPRKLRGALATLNQLFIVIGLASMFLIGAVVNWRTLALTGVAPCVVLFF   202
AtERDL16        SYVVPVYIAEISPKNLRGGLTTLNQLMIVIGSSVSFLIGSLISWKTLALTGLAPCIVLLF   211
AtERDL13        SYVVPVYIVEIAPKKVRGTFSAINSLVMCASVAVTYLLGSVISWQKLALISTVPCVFEFV   220
AtERDL1         SYVVPVYIAEITPKAFRGGFSFSNQLLQSFGISLMFFTGNFFHWRTLALLSAIPCGIQMI   194
AtERDL2         SYVVPVYIAEITPKTFRGGFSYSNQLLQCLGISLMFFTGNFFHWRTLALLSAIPSAFQVI   192
AtERDL9         SYVVPVYIAEITPKTVRGTFTFSNQLLQNCGVATAYYLGNFMSWRIIALIGILPCLIQLV   194
AtERDL12        SYVVPVYIAEITPKHVRGAFTFSNQLLQNCGVAVVYYFGNFLSWRTLAIIGSIPCWIQVI   193
AtERDL10        SYVVPVYIAEITPKHVRGAFSASTLLLQNSGISLIYFFGTVINWRVLAVIGALPCFIPVI   188
AtERDL11        SYVVPVYIAEITPKHVRGAFTASNQLLQNSGVSLIYFFGTVINWRVMAVIGAIPCILQTI   197
AtSFP1          SYVVPVYIAEITPKHVRGTFTFSNQLLQNAGLAMIYFCGNFITWRTLALLGALPCFIQVI   198
AtSFP2          SYVVPVYIAEISPKHVRGTFTFTNQLLQNSGLAMVYFSGNFLNWRILALLGALPCFIQVI   202
AtERD6          SYVIPVYIAEIAPKHVRGSFVFANQLMQNCGISLFFIIGNFIPWRLLTVVGLVPCVFHVF   225
AtERDL3         SYVIPVYIAEITPKHVRGAFVFANQLMQSCGLSLFYVIGNFVHWRNLALIGLIPCALQVV   199
                 :  *::* *::*: .**       :    .    :  *  . *: ::: .  *.

AtERDL14        LLFFIPESPRWLAKVGREKEVEGVLLSLRGAKSDVSDEAATILEYTKHVE-QQDIDSRGF   262
AtERDL15        LLFFIPESPRWLAKVGREMEVEAVLLSLRGEKSDVSDEAAEILEYTEHVKQQQDIDDRGF   250
AtERDL4         GLFFIPESPRWLAKMGLTDDFETSLQVLRGFETDITVEVNEIKRSVASSSK---RSAVRF   273
AtERDL6         GLFFIPESPRWLAKMGMTDEFETSLQVLRGFETDITVEVNEIKRSVASSTK---RNTVRF   272
AtERDL5         GLFVIPESPRWLAKVGKWEEFEIALQRLRGESADISYESNEIKDYTRRLTD---LSEGSI   256
AtERDL7         GLFFIPESPRWLAKVGRDTEFEAALRKLRGKKADISEEAAEIQDYIETLER---LPKAKM   252
AtERDL8         GTWFIPESPRWLEMVGRHSDFEIALQKLRGPQANITREAGEIQEYLASLAH---LPKATL   259
AtERDL16        GLCFIPESPRWLAKAGHEKEFRVALQKLRGKDADITNEADGIQVSIQALEI---LPKARI   268
AtERDL13        GLFFIPESPRWLSRNGRVKESEVSLQRLRGNNTDITKEAAEIKKYMDNLQE---FKEDGF   277
AtERDL1         CLFFIPESPRWLAMYGRERELEVTLKRLRGENGDILEEAAEIRETVETSRR---ESRSGL   251
AtERDL2         CLFFIPESPRWLAMYGQDQELEVSLKKLRGENSDILKEAAEIRETVEISRK---ESQSGI   249
AtERDL9         GLFFVPESPRWLAKEGRDEECEVVLQKLRGDEADIVKETQEILISVEAS------ANISM   248
AtERDL12        GLFFIPESPRWLAKKGRDKECEEVLQKLRGRKYDIVPEACEIKISVEASKK---NSNINI   250
AtERDL10        GIYFIPESPRWLAKIGSVKEVENSLHRLRGKDADVSDEAAEIQVMTKMLEE---DSKSSF   245
AtERDL11        GIFFIPESPRWLAKIRLSKEVESSLHRLRGKDTDVSGEAAEIQVMTKMLEE---DSKSSF   254
AtSFP1          GLFFVPESPRWLAKVGSDKELENSLFRLRGRDADISREASEIQVMTKMVEN---DSKSSF   255
AtSFP2          GLFFVPESPRWLAKVGSDKELENSLLRLRGGNADISREASDIEVMTKMVEN---DSKSSF   259
AtERD6          CLFFIPESPRWLAKLGRDKECRSSLQRLRGSDVDISREANTIRDTIDMTEN---GGETKM   282
AtERDL3         TLFFIPESPRLLGKWGHEKECRASLQSLRGDDADISEEANTIKETMILFDE---GPKSRV   256
                   .:***** *       :  .  *  *** . ::   *   *                 .

AtERDL14        FKLFQRKYALPLTIGVVLISMPQLGGLNGYTFYTDTIFTSTGVSS-DIGFILTSIVQMTG   321
AtERDL15        FKLFQRKYAFSLTIGVVLIALPQLGGLNGYSFYTDSIFISTGVSS-DFGFISTSVVQMFG   309
AtERDL4         VDLKRRRYYFPLMVGIGLLALQQLGGINGVLFYSSTIFESAGVTSSNVATFGVGVVQVVA   333
AtERDL6         VDLKRRRYYFPLMVGIGLLVLQQLGGINGVLFYSSTIFESAGVTSSNAATFGVGAIQVVA   332
AtERDL5         VDLFQPQYAKSLVVGVGLMVLQQFGGVNGIAFYASSIFESAGVSS-KIGMIAMVVVQIPM   315
AtERDL7         LDLFQRRYIRSVLIAFGLMVFQQFGGINGICFYTSSIFEQAGFPT-RLGMIIYAVLQVVI   311
AtERDL8         MDLIDKKNIRFVIVGVGLMFFQQFVGINGVIFYAQQIFVSAGASP-TLGSILYSIEQVVL   318
AtERDL16        QDLVSKKYGRSVIIGVSLMVFQQFVGINGIGFYASETFVKAGFTSGKLGTIAIACVQVPI   328
AtERDL13        FDLFNPRYSRVVTVGIGLLVLQQLGGLSGYTFYLSSIFKKSGFPN-NVGVMMASVVQSVT   336
AtERDL1         KDLFNMKNAHPLIIGLGLMLLQQFCGSSAISAYAARIFDTAGFPS-DIGTSILAVILVPQ   310
AtERDL2         RDLFHIGNAHSLIIGLGLMLLQQFCGSAAISAYAARIFDKAGFPS-DIGTTILAVILIPQ   308
AtERDL9         RSLFKKKYTHQLTIGIGLMLLQQLSGSAGLGYYTGSVFDLAGFPS-RIGMTVLSIVVVPK   307
AtERDL12        RSLFEKRYAHQLTIGIGLMLLQQLCGTAGISSYGSTLFKLAGFPA-RIGMMVLSLIVVPK   309
AtERDL10        CDMFQKKYRRTLVVGIGLMLIQQLSGASGITYYSNAIFRKAGFSE-RLGSMIFGVFVIPK   304
AtERDL11        SDMFQKKYRRTLVVGIGLMLIQQLSGASGITYYSNAIFRKAGFSE-RLGSMIFGVFVIPK   313
AtSFP1          SDLFQRKYRYTLVVGIGLMLIQQFSGSAAVISYASTIFRKAGFSV-AIGTTMLGIFVIPK   314
AtSFP2          CDLFQRKYRYTLVVGIGLMLIQQFSGSSAVLSYASTILRKAGFSV-TIGSTLLGLFMIPK   318
AtERD6          SELFQRRYAYPLIIGVGLMFLQQLCGSSGVTYYASSLFNKGGFPS-AIGTSVIATIMVPK   341
AtERDL3         MDLFQRRYAPSVVIGVGLMLLQQLSGSSGLMYYVGSVFDKGGFPS-SIGSMILAVIMIPK   315
                 .:        : :.. *: : *: *  .   *    :   *      .

AtERDL14        GVLGV-LLVDISGRRSLLLFSQAGMFLGCLATAISFFLQKNNC----WETGTPIMALISV   376
AtERDL15        GILGT-VLVDVSGRRFSSWNV-LGL--SYHSHFILLEGMENHC----WETGTPVLALFSV   361
AtERDL4         TGIAT-WLVDKAGRRLLLMISSIGMTISLVIVAVAFYLKEFVSPDSNMYNILSMVSVVGV   392
AtERDL6         TAIST-WLVDKAGRRLLLTISSVGMTISLVIVAAAFYLKEFVSPDSDMYSWLSILSVVGV   391
```

```
AtERDL5       TTLGV-LLMDKSGRRPLLLISATGTCIGCFLVGLSFSLQFVKQ----LSGDASYLALTGV   370
AtERDL7       TALNA-PIVDRAGRKPLLLVSATGLVIGCLIAAVSFYLKVHDM----AHEAVPVLAVVGI   366
AtERDL8       TALGATLLIDRLGRRPLLMASAVGMLIGCLLIGNSFLLKAHGL----ALDIIPALAVSGV   374
AtERDL16      TVLG-TILIDKSGRRPLIMISAGGIFLGCILTGTSFLLKGQSL----LLEWVPSLAVGGV   383
AtERDL13      SVLGI-VIVDKYGRRSLLTVATIMMCLGSLITGLSFLFQSYGL----LEHYTPISTFMGV   391
AtERDL1       SIIVM-FAVDRCGRRPLLMSSSIGLCICSFLIGLSYYLQNHGD----FQEFCSPILIVGL   365
AtERDL2       SIVVM-LTVDRWGRRPLLMISSIGMCICSFFIGLSYYLQKNGE----FQKLCSVMLIVGL   363
AtERDL9       AILGL-ILVERWGRRPLLMVM---------------------------------------   327
AtERDL12      SLMGL-ILVDRWGRRPLLMTSALGLCLSCITLAVAFGVKDVPG----IGKITPIFCFIGI   364
AtERDL10      ALVGL-ILVDRWGRRPLLLASAVGMSIGSLLIGVSFTLQEMNL----FPEFIPVFVFINI   359
AtERDL11      ALVGL-ILVDRWGRRPLLLASAVGMSIGSLLIGVSFTLQQMNV----LPELIPIFVFVNI   368
AtSFP1        AMIGL-ILVDKWGRRPLLMTSAFGMSMTCMLLGVAFTLQKMQL----LSELTPILSFICV   369
AtSFP2        AMIGV-ILVDKWGRRPLLLTSVSGMCITSMLIGVAFTLQKMQL----LPELTPVFTFICV   373
AtERD6        AMLAT-VLVDKMGRRTLLMASCSAMGLSALLLSVSYGFQSFGI----LPELTPIFTCIGV   396
AtERDL3       ALLGL-ILVEKMGRRPLLLASTGGMCFFSLLLSFSFCFRSYGM----LDELTPIFTCIGV   370
                :     ::  **:

AtERDL14      MVYFGSYGLGMGPIPWIIASEIYPVDVKGAAGTVCNLVTSISSWLVTYSFNFLLQWSSTG   436
AtERDL15      MVYFGSYGSGMGSIPWIIASEIYPVDVKGAAGTMCNLVSSISAWLVAYSFSYLLQWSSTG   421
AtERDL4       VAMVISCSLGMGPIPWLIMSEILPVNIKGLAGSIATLLNWFVSWLVTMTANMLLAWSSGG   452
AtERDL6       VAMVVFFSLGMGPIPWLIMSEILPVNIKGLAGSIATLANWFFSWLITMTANLLLAWSSGG   451
AtERDL5       LVYTGSFSLGMGGIPWVIMSEIFPIDIKGSAGSLVTVVSWVGSWIISFTFNFLMNWNPAG   430
AtERDL7       MVYIGSFSAGMGAMPWVVMSEIFPINIKGVAGGMATLVNWFGAWAVSYTFNFLMSWSSYG   426
AtERDL8       LVYIGSFSIGMGAIPWVIMSEIFPINLKGTAGGLVTVVNWLSSWLVSFTFNFLMIWSPHG   434
AtERDL16      LIYVAAFSIGMGPVPWVIMSEIFPINVKGIAGSLVVLVNWSGAWAVSYTFNFLMSWSSPG   443
AtERDL13      LVFLTSITIGIGGIPWVMISEMTPINIKGSAGTLCNLTSWSSNWFVSYTFNFLFQWSSSG   451
AtERDL1       VGYVLSFGIGLGGLPWVIMSEVFPVNVKITAGSLVTVSNWFFSWIIIFSFNFMMQWSAFG   425
AtERDL2       VGYVSSFGIGLGGLPWVIMSEIFPVNVKITAGSLVTMSNWFFNWIIIYSFNFMIQWSASG   423
AtERDL9       ------------------------------------------------------------   327
AtERDL12      LSFTMMFAIGMGALPWIIMSEIFPMDIKVLAGSLVTIANWFTGWIANYAFNFMLVWSPSG   424
AtERDL10      LVYFGFFAIGIGGLPWIIMSEIFPINIKVSAGSIVALTSWTTGWFVSYGFNFMFEWSAQG   419
AtERDL11      LVYFGCFAFGIGGLPWVIMSEIFPINIKVSAGTIVALTSWTSGWFVSYAFNFMFEWSAQG   428
AtSFP1        MMYIATYAIGLGGLPWVIMSEIFPINIKVTAGSIVTLVSFSSSSIVTYAFNFLFEWSTQG   429
AtSFP2        TLYIGTYAIGLGGLPWVIMSEIFPMNIKVTAGSIVTLVSWSSSSIVTYAFNFLLEWSTQG   433
AtERD6        LGHIVSFAMGMGGLPWIIMAEIFPMNVKVSAGTLVTVTNWLFGWIITYTFNFMLEWNASG   456
AtERDL3       VGFISSFAVGMGGLPWIIMSEIFPMNVKVSAGTLVTLANWSFGWIVAFAYNFMLEWNASG   430


AtERDL14      TFMMFATVMGLGFVFTAKLVPETKGKSLEEIQSAFTDSTSEDSTIF         482
AtERDL15      TFLMFATVAGLGFVFIAKLVPETKGKSLEEIQSLFTDSPPQDSTIF         467
AtERDL4       TFTLYALVCGFTVVFVSLWVPETKGKTLEEIQALFR----------         488
AtERDL6       TFTLYGLVCAFTVVFVTLWVPETKGKTLEELQSLFR----------         487
AtERDL5       TFYVFATVCGATVIFVAKLVPETKGRTLEEIQYSIGYVEL------         470
AtERDL7       TFLIYAAINALAIVFVIAIVPETKGKTLEQIQAIVNP---------         463
AtERDL8       TFYVYGGVCVLAIIFIAKLVPETKGRTLEEIQAMMM----------         470
AtERDL16      TFYLYSAFAAATIIFVAKMVPETKGKTLEEIQACIRRET-------         482
AtERDL13      VFFIYTMISGVGILFVMKMVPETRGRSLEEIQAAITR---------         488
AtERDL1       TYFIFAGVSLMSFVFVWTLVPETKGRTLEDIQQSLGQLS-------         464
AtERDL2       TYFIFSGVSLVTIVFIWTLVPETKGRTLEEIQTSLVRLS-------         462
AtERDL9       ----------------------------------------------         327
AtERDL12      TFIISAIICGATIVFTWCLVPETRRLTLEEIQLSFVNV--------         462
AtERDL10      TFYIFAMVGGLSLLFIWMLVPETKGQSLEELQASLTGTT-------         458
AtERDL11      TFYIFAAVGGMSFIFIWMLVPETKGQSLEELQASLTGTS-------         467
AtSFP1        TFFIFAGIGGAALLFIWLLVPETKGLSLEEIQVSLIHQPDERNQT-         474
AtSFP2        TFYVFGAVGGLALLFIWLLVPETKGLSLEEIQASLIREPDRINQS-         478
AtERD6        MFLIFSMVSASSIVFIYFLVPETKGRSLEEIQALLNNSVQ------         496
AtERDL3       TFLIFFTICGAGIVFIYAMVPETKGRTLEDIQASLTDFLQ------         470
```
